# Feeder-Free Generation of Endocardial and Cardiac Valve Cells from Human Pluripotent Stem Cells

**DOI:** 10.1101/2023.03.03.530824

**Authors:** Clifford Z. Liu, Aditi Prasad, Bharati Jadhav, Andrew J. Sharp, Bruce D. Gelb

**Affiliations:** Mindich Child Health and Development Institute, Icahn School of Medicine at Mount Sinai, New York, NY, USA; Department of Cell, Developmental, and Regenerative Biology, Icahn School of Medicine at Mount Sinai, New York, NY, USA; Department of Pediatrics, Icahn School of Medicine at Mount Sinai, New York, NY, USA; Department of Genetics and Genomic Sciences, Icahn School of Medicine at Mount Sinai, New York, NY, USA

**Author notes:** Author to whom correspondence should be addressed: Bruce D. Gelb, MD Icahn School of Medicine at Mount Sinai One Gustave Levy Place, Box 1040 New York, NY 10029.

## Abstract

Valvular heart disease presents a significant health burden, yet advancements in valve biology and novel therapeutics have been hindered by the lack of accessibility to human valve cells. In this study, we have developed a scalable and feeder-free method to differentiate human induced pluripotent stem cells (iPSCs) into endocardial cells. Importantly, we show that these endocardial cells are transcriptionally and phenotypically distinct from vascular endothelial cells and can be directed to undergo endothelial-to-mesenchymal transition (EndMT) to generate cardiac valve cell populations. Following this, we identified two distinct populations—one population undergoes EndMT to become valvular interstitial cells (VICs), while the other population reinforces their endothelial identity to become valvular endothelial cells (VECs). Lastly, we confirmed the identities of our iPSC-derived cell populations and identified putative markers through transcriptomic analyses. By increasing the accessibility to these cell populations, we aim to accelerate discoveries for cardiac valve biology and disease.

## Introduction

Valvular heart disease represents a significant health burden, with an estimated total U.S. prevalence of 2.5%.^1^ Additionally, aberrant development of cardiac valves is a frequent occurrence in congenital heart disease, representing 20-30% of cases.^2^ Despite this high prevalence, valvular disease is primarily managed through surgical intervention, with limited pharmacological therapies available.^3, 4^ A major barrier towards the development of targeted therapeutics for valvular disease is the lack of appropriate *in vitro* systems to model both developmental and disease processes with human cells. One method of circumventing this barrier would be to develop a robust and scalable differentiation strategy by directing pluripotent stem cells (PSCs) towards normal cardiac valve developmental pathways.

Cardiac valvulogenesis is a complex process that involves numerous cell types and signaling pathways (reviewed extensively elsewhere, see ^2, 5, 6^). Briefly, cardiac valvulogenesis begins after looping of the primitive heart tube, during which BMP signaling from the myocardium leads to increased extracellular matrix (ECM) deposition and swelling of the cardiac jelly at the atrioventricular canal and outflow tract.^5–8^ A subpopulation of the endocardial cells overlaying the cardiac jelly then undergoes endothelial-to-mesenchymal transition (EndMT) to become the putative valvular interstitial cells (VICs) and migrates into the cardiac jelly to populate the developing endocardial cushion.^8^ Multiple signaling pathways have been implicated in promoting EndMT, including BMP2/4,^7, 9^ TGF*β*2,^10^ NOTCH,^11, 12^ WNT/*β*-CATENIN,^13^ and HIPPO/YAP.^14^ Following EndMT, the endocardial cushions begin to elongate into valve leaflets—a process resulting from proliferation, apoptosis, and the secretion and remodeling of ECM proteins by the VICs.^15–17^ Notably, a subpopulation of endocardial cells escapes EndMT and continues to overlay the endocardial cushion to become valvular endothelial cells (VECs).^6^ Although other lineages, such as neural crest derivatives,^18, 19^ epicardial cells,^20–22^ and hemogenic populations,^23, 24^ also contribute to valve development, these are not the focus of the present study.

Given the central role of endocardium in valve development, generating an endocardial population from PSCs is of intense interest. Some previous studies have utilized VEGF to direct PSCs into an endothelial population, transitioning through a cardiac mesoderm stage^25–28^ or epicardial intermediate.^29^ Intriguingly, these endothelial populations express some endocardial markers and, in some cases, have the potential to undergo EndMT.^25, 28^ However, recent work by Mikryukov *et al*. demonstrated that endocardial cells could be differentiated without the use of VEGF but rather via a combination of BMP10 and bFGF.^30^ In this strategy, PSCs were first maintained on mouse embryonic fibroblast feeder cells and then formed into embryoid bodies (EBs). These EBs were then directed towards a mesoderm fate with BMP4, ACTIVIN A, and bFGF prior to dissociation and endocardial specification with BMP10 and bFGF. Critically, this group demonstrated that this BMP10-induced endocardial population is transcriptionally and functionally distinct from VEGF-induced endothelial populations, has robust expression of endocardial markers, and maintains EndMT potential.

Feeder cells, such as mouse embryonic fibroblasts, are routinely used in PSC cultures to help support their pluripotency^31, 32^ and have also been utilized to support a VEGF-directed strategy of differentiating endocardial-like cells.^25^ However, there are specific disadvantages to the use of feeder cells. In addition to increasing the technical difficulty of culturing PSCs, feeder cells are known to secrete various growth factors and cytokines, hindering the ability to precisely control the levels of signaling molecules that PSCs encounter. Incomplete removal and presence of residual feeder cells has also been shown to influence transcriptomic analysis of PSCs and can limit the suitability of differentiated cells for clinical purposes.^33^

To circumvent these limitations, we sought to develop a feeder-free, monolayer differentiation strategy incorporating GSK3 inhibition with CHIR-99021 (CHIR) that would be compatible with the BMP10-directed endocardial specification strategy. Since the discovery of efficient cardiomyocyte differentiation by GSK3 inhibition,^34, 35^ CHIR has been widely used for monolayer differentiation into mesoderm derivatives, including cardiomyocytes and vascular endothelium.^36, 37^ Utilizing a small molecule for such differentiations not only allows for increased scalability and adaptability but can also decrease overall costs.^38^ In this study, we utilized two wild-type human induced pluripotent stem cell (iPSC) lines to demonstrate that endocardial cells can be generated via CHIR in a feeder-free monolayer system. We confirmed that BMP10-induced endocardial cells are distinct from VEGF-induced endothelial cells and that these endocardial cells maintain their ability to undergo EndMT. Furthermore, we showed that following an EndMT challenge, two distinct populations emerge and are representative of VICs and VECs. Lastly, we performed bulk RNA sequencing on these populations and identified putative markers to distinguish them. The ability to generate these populations in a scalable fashion will facilitate novel studies into human valvulogenesis and valvular disease.

## Results

### Monolayer differentiation of feeder-free iPSCs into endocardium

Since the identification of the utility of GSK3 inhibition in generating cardiomyocytes,^34, 35^ the use of CHIR has been widely adapted in differentiating various mesoderm derivatives. To determine if this strategy would be compatible with producing endocardium, we utilized two wild-type human iPSC lines: WTC11 and MSN02-4. These iPSCs were grown on Matrigel without the use of feeder cells and subsequently directed to a mesoderm fate with 48 h of CHIR (8 μM). Afterwards, we introduced BMP10 (10 ng/mL) and bFGF (50 ng/mL) over the next 12 days to generate a CD31^+^ (PECAM1) endocardial population. As a control, we also differentiated iPSCs towards a vascular endothelial fate with VEGF-A (100 ng/mL) and bFGF (50 ng/mL) (Figure 1A).

**Figure 1:**
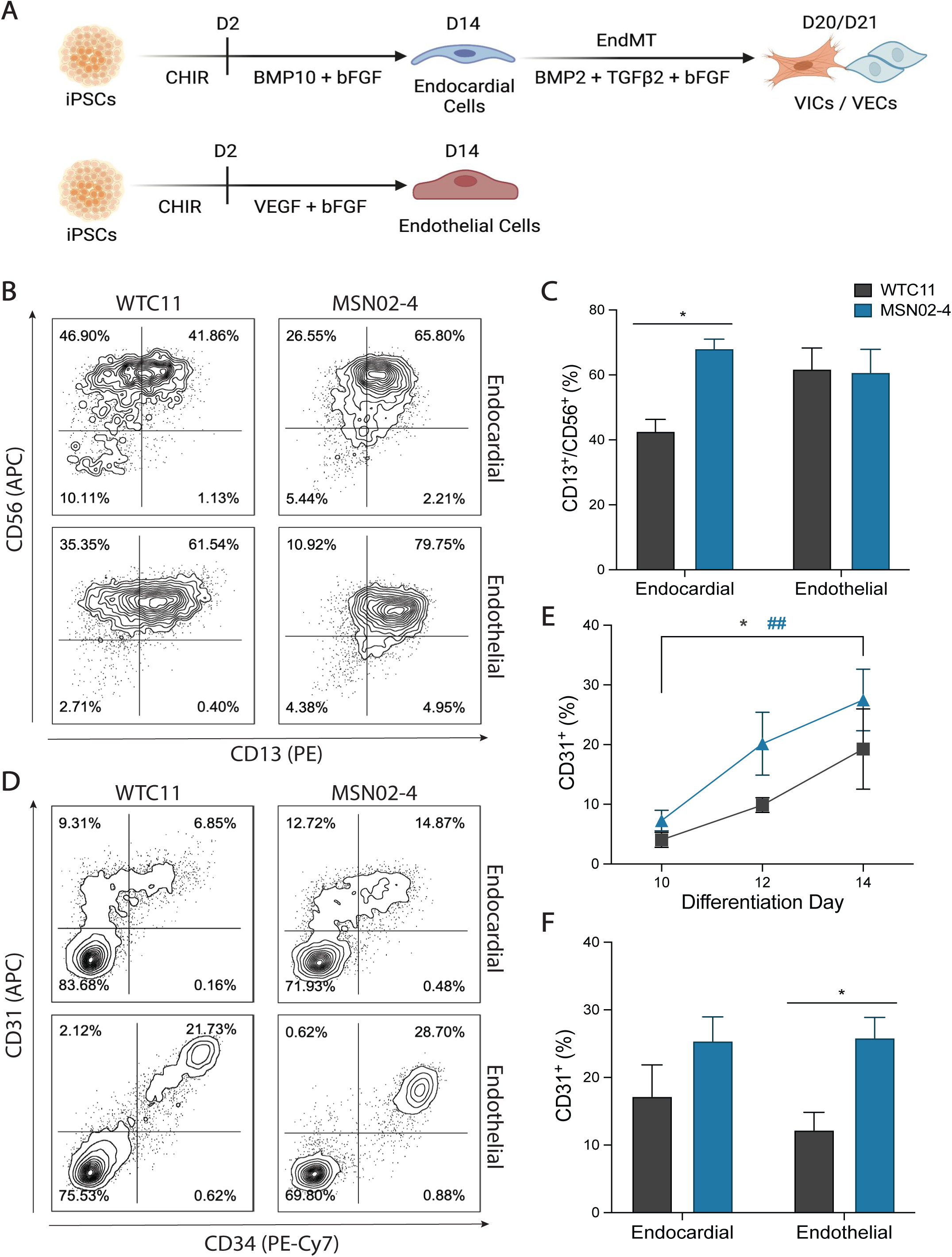
Feeder-free generation of endocardial and endothelial populations from iPSCs. **(A)** Schematic of differentiation timeline. **(B)** Flow cytometry plots assessing mesoderm generation on Day 4 of their respective differentiation strategies. **(C)** Quantification of CD13^+^/CD56^+^ mesoderm generation (n = 4, *p < 0.05, 2-way ANOVA with Sidak’s multiple comparisons test, error bars ± SEM). **(D)** Flow cytometry plots assessing endocardial/endothelial differentiation efficiency on Day 14. **(E)** Timeline of CD31^+^ endocardial differentiation on Days 10, 12, and 14 (n = 4, *p < 0.05 for WTC11, ##p < 0.01 for MSN02-4, 2-way ANOVA with Dunnett’s multiple comparisons test, error bars ± SEM). **(F)** Quantification of CD31^+^ endocardial efficiency on Day 14 (n = 5, *p < 0.05, 2way ANOVA with Sidak’s multiple comparisons test, error bars ± SEM).

To determine the efficiency of mesoderm generation for each of the lines, we then dissociated differentiating iPSCs on Day 4 and evaluated them for co-expression of CD13 and CD56 (Figure 1B-C). As CD56 has been identified as a marker of early mesoderm-committed populations^39^ and CD13 as an early pre-cardiac mesoderm marker,^40^ CD13^+^/CD56^+^ double-positive cells represent a mesoderm population capable of undergoing further cardiac cell-type specification. We found that both the endocardial and control endothelial differentiation conditions generated a robust CD13^+^/CD56^+^ population in the MSN02-4 line (∼60% yield). However, the WTC11 line had diminished CD13^+^/CD56^+^ specification under the endocardial protocol (∼40% yield), while the endothelial protocol performed similarly with these iPSCs as did the MSN02-4 line (∼60% yield).

We then assessed the induction of endocardial cells on Days 10, 12, and 14 (Figure 1E). Over these timepoints, both the WTC11 and MSN02-4 lines demonstrated increased specification of an endocardial population, marked by cell surface expression of CD31. By Day 14, both iPSC lines produced an endocardial population, with yield ranging from 15-30% of the entire population (Figure 1D-F). Additionally, we found that the endocardial population had heterogenous expression of CD34, indicative of endocardial cells at varying levels of maturation. When we directed our iPSCs towards an endothelial population with VEGF-A, we generated a distinct CD31^+^/CD34^+^ population by Day 14. Interestingly, the WTC11 line appeared to be less efficient at generating this endothelial population compared to the MSN02-4 line (∼12% vs. ∼25% yield).

### BMP10-derived endocardial cells upregulate endocardial-specific genes

Following specification of both endocardial and endothelial cell populations on Day 14, these populations were isolated using magnetic-activated cell sorting (MACS) for CD31 and CD34, respectively. This strategy allowed us to routinely isolate these populations to ∼95% or greater purity (Figure S1A-B). We then performed RT-qPCR to assess the expression of cardiac- and endocardial-specific genes (Figure 2A). Crucially, we found that for both the WTC11 and MSN02-4 lines, BMP10-derived endocardial cells upregulated expression of cardiac-specific transcription factors *NKX2.5*, *GATA4*, and *GATA5* compared to the endothelial population. In addition, we found upregulation of endocardial genes *ISL1, HAPLN1, NPR3, NRG1, MEIS2,* and *TMEM100* in our endocardial population. Interestingly, *NFATc1* expression was detected at similar levels between both endocardial and endothelial populations.

**Figure 2:**
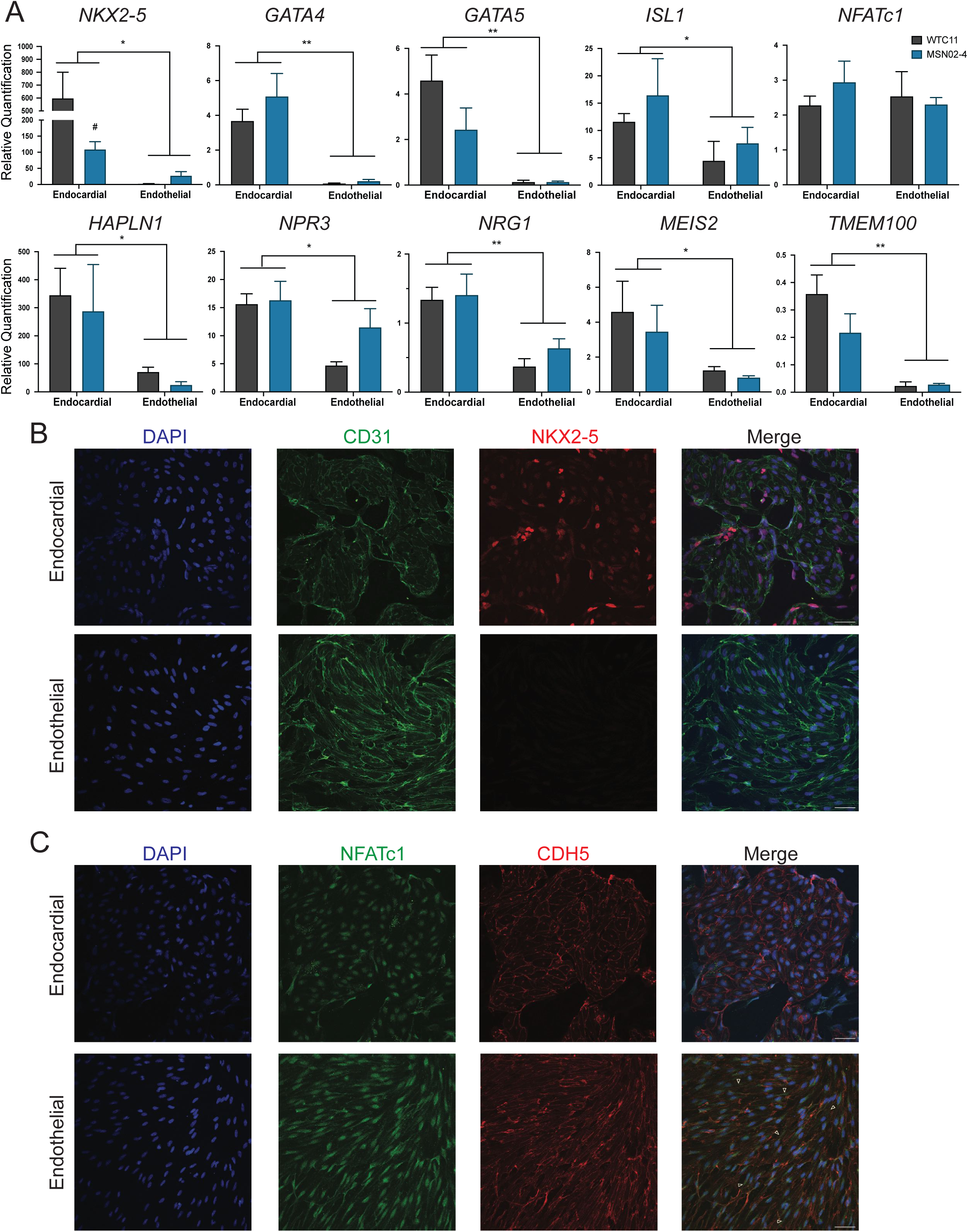
BMP10-derived endocardial cells are distinct from VEGF-derived endothelial cells. **(A)** RT-qPCR analysis of cardiac and endocardial gene expression on Day 14 endocardial and endothelial cells (n = 4, *p < 0.05, **p < 0.01, 2-way ANOVA, error bars ± SEM). **(B)** Confocal microscopy of endocardial and endothelial cells stained for CD31 (GFP) and NKX2-5 (RFP). Scale bar = 50 μm. **(C)** Confocal microscopy of endocardial and endothelial cells stained for NFATc1 (GFP) and CDH5 (RFP). White arrowheads = diffuse NFATc1 in the cytoplasm of endothelial cells. Scale bar = 50 μm.

Next, we performed confocal microscopy on endocardial and endothelial cells. Both endocardial and endothelial populations had expression of CD31, localized to the plasma membrane. Notably, only endocardial cells had co-expression of CD31 and NKX2.5, while NKX2.5 was undetectable in the endothelial cells (Figure 2B). Additionally, although we detected NFATc1 in both the endocardial and endothelial cells, we found that NFATc1 was predominantly localized in the nucleus in endocardial cells while the NFATc1 staining pattern was more diffuse in the endothelial cells (Figure 2C). Lastly, we showed that both the endocardial and endothelial populations co-localized CD31 and CDH5 (VE-CADHERIN) to the plasma membrane (Figure S2).

### EndMT challenge of endocardial cells generates valvular interstitial cells and valvular endothelial cells

After sorting out the endocardial population on Day 14 with MACS, we then plated the endocardial cells on fibronectin and induced EndMT with a combination of BMP2 (100 ng/mL), TGF*β*2 (0.3 ng/mL), and bFGF (10 ng/mL) (Figure 1A). Following 6-7 days of this EndMT condition, we found that a subpopulation of the endocardial cells lost CD31 expression to become VICs and began to express the mesenchymal surface marker PDGFR*β*. Additionally, there was a population that escaped the EndMT process and maintained CD31 expression, becoming VECs (Figure 3A-B). Interestingly, we found that our endothelial population was unable to survive this EndMT process, with nearly 100% cell death observed by the end of the EndMT period (data not shown).

**Figure 3:**
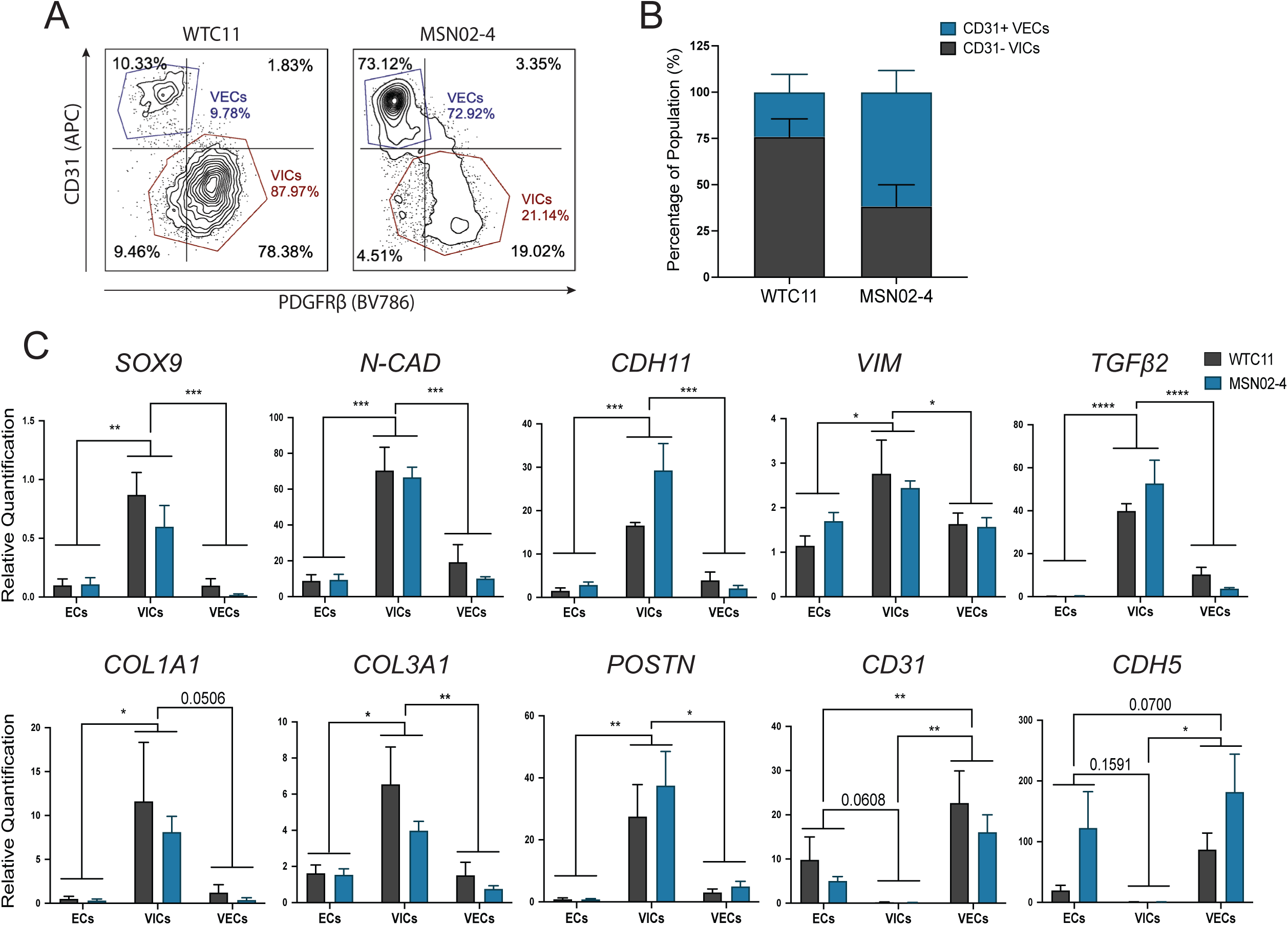
EndMT of endocardial cells generates VIC and VEC populations. **(A)** Flow cytometry plots following EndMT. **(B)** Quantification of CD31^-^ VIC and CD31^+^ VEC populations following EndMT (n = 4, error bars ± SEM). **(C)** RT-qPCR analysis of mesenchymal and endothelial gene expression on Day 14 endocardial cells, VICs, and VECs (n = 4, *p < 0.05, **p < 0.01, ***p < 0.001, ****p < 0.0001, 2-way ANOVA with Tukey’s multiple comparisons test, error bars ± SEM).

By utilizing our CD31 MACS strategy, we were then able to sort the corresponding CD31^-^ VICs from CD31^+^ VECs (Figure S1C-D). RT-qPCR analysis on the starting Day 14 endocardial cells and the post-EndMT VIC/VEC populations revealed that VICs strongly upregulated mesenchymal-specific genes, such as *SOX9, N-CAD, CDH11, VIM, TGFβ2, COL1A1, COL3A1,* and *POSTN*. Furthermore, the VICs lost expression of endothelial genes *CD31* and *CDH5* (Figure 3C). Interestingly, we found that, following the EndMT challenge, VECs reinforced their endothelial identity and upregulated endothelial genes *PECAM1* and *CDH5* compared to Day 14 endocardial cells.

Next, we performed confocal microscopy on both VICs and VECs. Notably, we found that VICs became morphologically distinct from their initial endocardial population; in addition to having larger cell sizes, these cells displayed a more elongated morphology (Figure 4A-B). Furthermore, we found that the VICs had robust staining of mesenchymal markers PERIOSTIN, VIMENTIN, *α*SMA, and SOX9. Different from the morphology of the VICs, the VECs exhibited typical cuboidal endothelial morphology and expressed endothelial proteins CD31 and CDH5 at the plasma membrane (Figure 4C).

**Figure 4:**
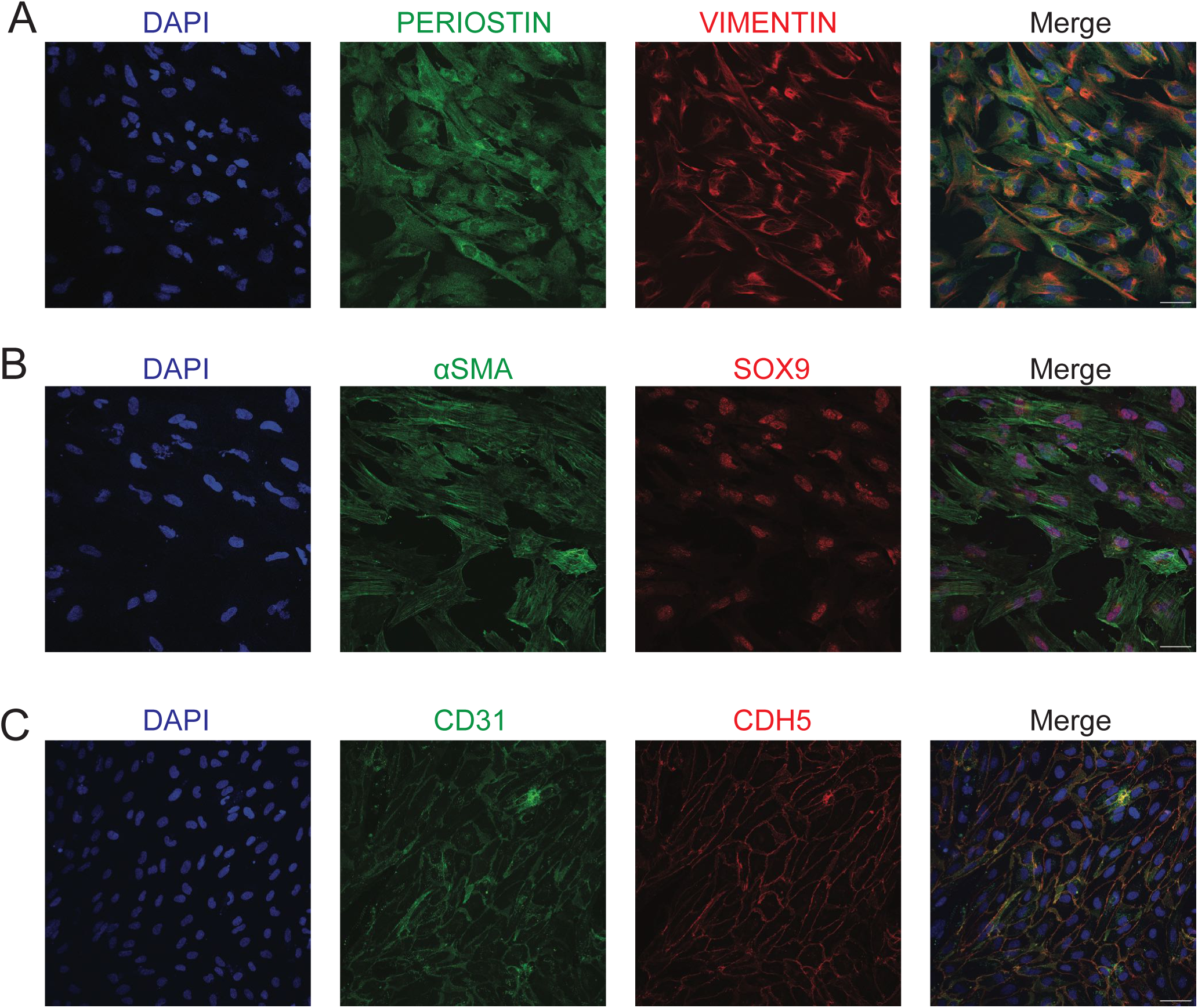
Confocal microscopy of VICs and VECs. **(A)** VICs stained for PERIOSTIN (GFP) and VIMENTIN (RFP). **(B)** VICs stained for *α*SMA (GFP) and SOX9 (RFP). **(C)** VECs stained for CDH5 (GFP) and CD31 (RFP).

### Transcriptional analysis of iPSC-derived endothelial cells, endocardial cells, VICs, and VECs

To further validate each of our respective cell types and identify potential markers, we conducted bulk RNA sequencing with cells from our two wild-type iPSC lines. Results from principal component analysis (PCA) showed that the two iPSC lines for each cell type grouped together, with the first two principal components (PC1 and PC2) accounting for 55% and 18% of the variance, respectively (Figure S3A). Importantly, VICs were the most distinct from the control endothelial cells, separated primarily by PC1. We also conducted a differential gene expression analysis and performed unsupervised hierarchical clustering on the upper quartile of genes ranked by variance. This revealed that our two genotypes for each cell type grouped together, with all endothelial cell types clustering together, and the two cardiac endothelial subtypes (endocardial and VECs) clustering closer than the control endothelial population (Figure S3B).

Next, we assessed the top differentially expressed genes (DEGs) between the endothelial and endocardial cell types. Notably, we found that genes upregulated in the endocardial cells include the cardiac-specific transcription factors *NKX2.5* and *GATA4* (Figure 5A-B). Some other genes that were strongly upregulated in the endocardial cells include *TNR, ANGPT1, CRHBP, CLEC4M, FGF23, NTRK2,* and *MRC1*. Gene ontology (GO) analysis of upregulated DEGs in the endocardial cells revealed extracellular matrix development terms as well as heart development, including ‘ventricular cardiac muscle tissue morphogenesis’ and ‘ventricular septum development’ (Figure 5C). Intriguingly, the 21^st^ GO term when ranked by p-values was ‘endocardial cushion development’ (p = 0.0005). Analysis of endothelial cell DEGs revealed upregulation of endothelial or blood vessel development genes, such as *CXCR4, PGF, ESM1, TNFSF10,* and *NOTCH4* (Figure 5A-B). Most significant GO terms resulting from upregulated endothelial DEGs included ‘endothelial cell development’ and ‘regulation of angiogenesis’ (Figure 5D).

**Figure 5:**
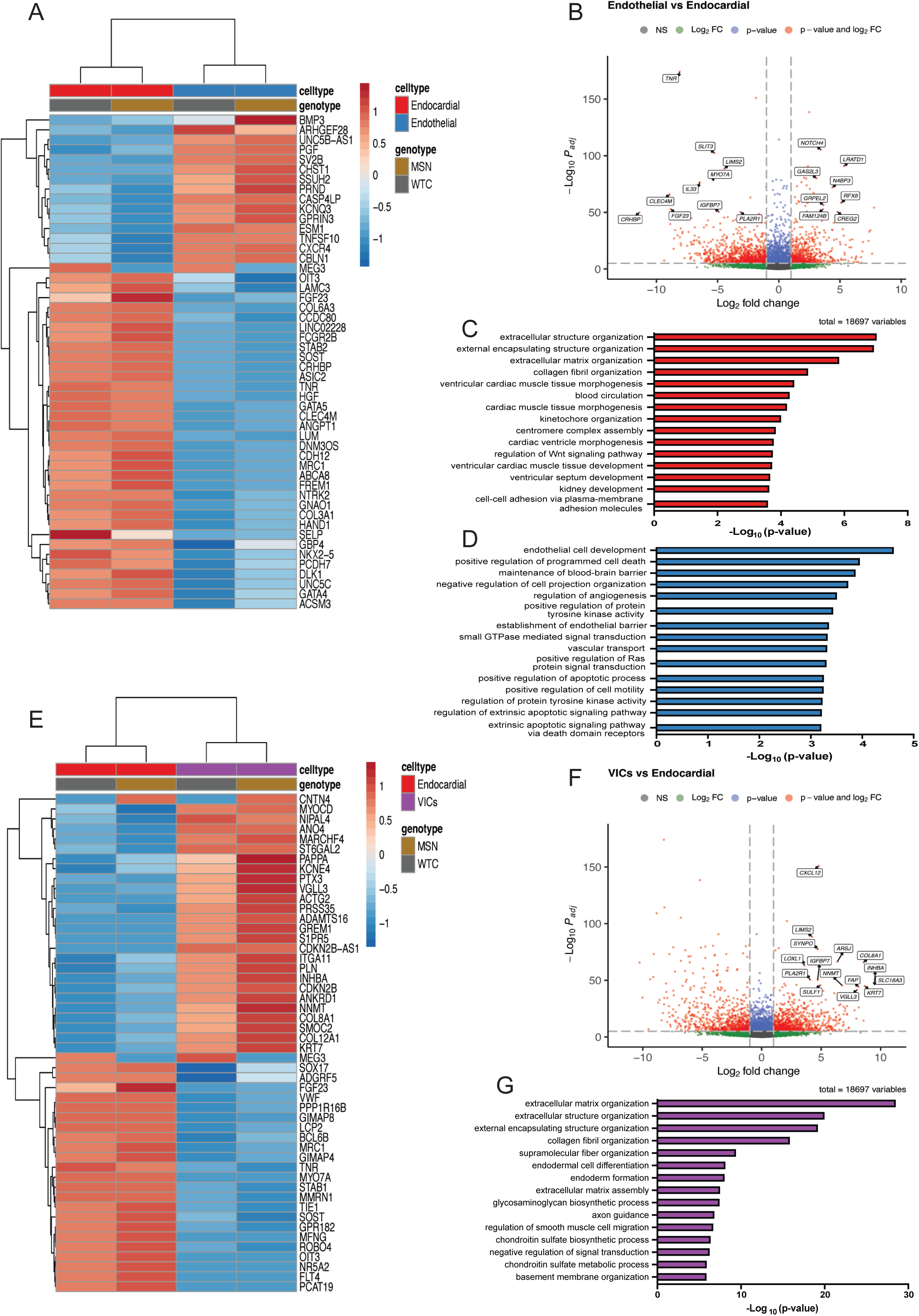
Bulk-RNAseq analysis of iPSC-derived populations. **(A)** Heatmap of the top 50 DEGs ranked by variance between endothelial and endocardial populations. **(B)** Volcano plot of DEGs between endothelial and endocardial populations. **(C)** Top 15 biological process GO terms for upregulated DEGs in endocardial cells, ranked by p-value. **(D)** Top 15 biological process GO terms for upregulated DEGs in endothelial cells, ranked by p-value. **(E)** Heatmap of top 50 DEGs ranked by variance between VIC and endocardial populations. **(F)** Volcano plot of DEGs between VIC and endocardial populations. **(G)** Top 15 biological process GO terms for upregulated DEGs in VICs, ranked by p-value.

Analysis of DEGs between VICs and endocardial cells revealed upregulation of extracellular matrix genes, such as *COL8A1, COL12A1, ADAMTS16,* and *ITGA11*. Additionally, we found significant upregulation of genes involved in valve development or VIC function, such as *CXCL12, LOXL1,* and *VGLL3* (Figure 5E-F). GO analysis on upregulated VIC DEGs revealed numerous terms involving the extracellular matrix, including ‘extracellular matrix organization’, ‘collagen fibril organization’, and ‘glycosaminoglycan biosynthetic process’ (Figure 5G).

We then assessed DEGs between VECs and endocardial cells and found numerous significantly upregulated genes, including *CXCL12, IL33, SIRPB2, CLDN5, CREG2, PALMD,* and *AQP1*. However, as VECs are very similar to endocardial and endothelial cells, we also assessed for genes uniquely upregulated in VECs compared to both endocardial and endothelial cells (Figure S4A-B). This analysis yielded many upregulated genes in the VECs, including *HES5, SAMD9L, INHBA, VGLL3,* and *COL8A1.* To confirm the endothelial identity of VECs and to increase the contrast between comparisons, we further generated DEGs between VECs and VICs (Figure S4C). Indeed, this comparison revealed upregulation of endothelial genes in the VECs, such as *SOX17, CDH5, TIE1, FLT4, CD34,* and *CXCR4*. GO term analysis of upregulated genes in VECs from this analysis revealed terms related to endothelial cell functions, including ‘regulation of angiogenesis’, ‘angiogenesis’, and ‘regulation of endothelial cell migration’ (Figure S4D). Additionally, we also analyzed BioPlanet GO terms to dissect differential signaling pathways between VICs and VECs (Figure S4E-F). We found that the most significant terms associated with VICs included ‘TGF-beta regulation of extracellular matrix’ and ‘Beta-1 integrin cell surface interactions’. The most significant term for the VECs, however, was ‘TNF-alpha effects on cytokine activity, cell motility, and apoptosis.’ Collectively, these data support the differentiation of iPSCs into an endocardial population with BMP10 that are distinct from VEGF-derived endothelial cells, and that these endocardial cells can then undergo an EndMT challenge to generate both VICs and VECs.

## Discussion

The ability for iPSCs to differentiate into various cell types has allowed researchers to further their understanding of development and the biological mechanisms that govern healthy and disease states. Expanding the repertoire of cell types that can be differentiated from iPSCs will only further these goals. In the present study, we demonstrated a novel protocol that allowed us to differentiate iPSCs into endocardial cells as a monolayer without requiring the presence feeder cells or the formation of embryoid bodies in suspension. These endocardial cells were then challenged to undergo EndMT, resulting in the segregation of two populations—VICs and VECs.

Our protocol was validated using two wild-type iPSC lines. We found that CHIR alone was sufficient to induce mesoderm specification in iPSCs cultured as a monolayer and that the efficiency of our mesoderm specification under the endocardial condition differed between our two wild-type iPSC lines (Figure 1C). Notably, we have found that this efficiency can be improved by titrating the CHIR concentration and that improving this step can increase the efficiency of endocardial generation by Day 14 (data not shown). These differences suggest that the genetic background can influence iPSC response to CHIR and that optimal CHIR concentrations should be determined for each potential cell line.

Following specification of mesoderm, we promoted endocardial differentiation with BMP10 and bFGF and found that key cardiac transcription factors, such as *NKX2.5, GATA4,* and *GATA5*, were uniquely upregulated in our endocardial population compared to VEGF-derived endothelial cells (Figure 2A). These results are in keeping with the findings from Mikryukov *et al.* that BMP10-derived endocardial cells are transcriptionally and phenotypically distinct from VEGF-derived endothelial cells.^30^ Interestingly, we found that NFATc1 was expressed in both of our endocardial and endothelial populations and that it was predominantly nuclear localized within the endocardial population while endothelial cells had NFATc1 diffusely throughout the cell (Figure 2C). This is consistent with prior work identifying NFATc1 as one of the earliest markers of the endocardium, as its expression in E8.5 mice is restricted to the heart tube endocardium and becomes specific to the cushion endocardium between E9.5 to E11.5.^41, 42^ In this context, NFATc1 is found in the nucleus and functions to regulate EndMT.^43^ In endothelial cells, however, NFATc1 expression has also been described, where it functions to regulate angiogenesis genes^44^ and is also involved in the development and patterning of lymphatic endothelial cells.^45^ This suggests that NFATc1 specificity to the endocardium is temporally restricted and that NFATc1 levels may be reduced or absent at earlier time points for our endothelial cells.

After EndMT of our endocardial cells, we identified two distinct populations that represent VICs and VECs. Multiple lines of evidence support the establishment of these cell types. First, our transcriptomic data confirmed the divergence of endocardial cells from endothelial cells, with significant GO terms involving both cardiac developmental processes, extracellular matrix organization, and specifically, ‘endocardial cushion development’ (Figure 5A-D). Given the normal developmental trajectory for valvulogenesis, the populations derived from EndMT of the endocardial cells should represent cardiac valve populations. Indeed, our transcriptional and immunofluorescence analyses confirmed that VICs are of a mesenchymal identity and are involved in extracellular matrix production and organization, including production of both collagen and glycosaminoglycans (Figures 3C, 4A-B, and 5E-G). As such, we detected upregulation of numerous extracellular matrix genes, such as *COL8A1, COL12A1,* and *ADAMTS16*, in addition to genes that are known to have a function in VICs, such as *CXCL12*^46^ and *LOXL1.*^47^ Furthermore, transcriptional and immunofluorescence analyses revealed that VECs reinforced their endothelial identity after being challenged with EndMT and that they upregulated gene related to GO terms involved with endothelial cell functions (Figures 3C, 4C, and S4). By comparing our transcriptomic data for VECs with endocardial and endothelial populations, we identified numerous genes uniquely upregulated in VECs (Figure S5). Some of these genes, such as *PALMD* and *AQP1,* have also been recently identified through single-cell RNA sequencing of embryonic mouse atrioventricular canals as putative VEC-specific markers.^48^

Our transcriptomic analysis also identified genes that were upregulated in both VICs and VECs, such as *CXCL12, VGLL3,* and *INHBA*, which may serve as transcriptional markers that distinguish these valve cells from other endothelial and mesenchymal populations. *CXCL12* is known to be important in valve development, as *Cxcl12*-null mice develop dysplastic and misaligned semilunar valves.^46^ CXCL12 signaling has also been shown to influence positioning and orientation of cells undergoing EndMT, along with regulation of VIC proliferation during remodeling and maturation of the valve leaflet.^46^ Additionally, upregulation of *VGLL3* is particularly intriguing as its function in valve development, to the best of our knowledge, is unexplored. However, in other contexts it is thought to activate HIPPO signaling and regulate YAP activity by competing for TEAD binding.^49^ Notably, the function of *Vgll4* has recently been explored in mice, where knockout of *Vgll4* led to semilunar valve thickening—a phenotype that was rescued with partial knockout of *Yap*.^50^ As YAP also responds differentially to changes in mechanical force and regulates valve growth and remodeling,^51^ it would be worth investigating the influence of VGLL3 on these processes and how its function differs from VGLL4.

In summary, our study shows that generating endocardial and valve cell populations from iPSCs as a monolayer is feasible and an easily scalable approach that overcomes technical challenges. As iPSC-derived cell populations are generally considered to be fetal-like, our protocol would be particularly advantageous for modeling congenital valve disease. Furthermore, the transcriptomic dataset we have generated can aid in identifying novel biology and potential markers for these cells. With the development of this protocol, we hope to expand access to these human cell populations and advance discoveries in cardiac valve biology and disease.

## Limitations

Our study was limited to bulk-RNAseq analysis, precluding the identification of subpopulations within each of cell types. To address this, future directions include performing single-cell RNA sequencing to explore the heterogeneity of our populations. Additionally, we only cultured our cells in 2D without the influence of mechanical forces. Future work can include the generation of tissues with these cells and examining the effects of mechanical stimuli on signaling between cell populations. Lastly, we have created a purely *in vitro* system and have not yet demonstrated how our cells compare to their respective populations *in vivo*.

## Supporting information

List of DEGs

## Acknowledgments

We extend our gratitude to Dr. Nicole Dubois for providing the MSN02-4 iPSC line and experimental advice as well as to the lab of Dr. Gordon Keller for fruitful discussions on our results. We also acknowledge the Flow Cytometry CORE, Microscopy CORE, and Scientific Computing and Data at the Icahn School of Medicine at Mount Sinai for their valuable support. This project was made possible by the generous funding from the National Institutes of Health/National Heart, Lung, and Blood Institute (R35 HL135742), the American Heart Association/The Children’s Heart Foundation Predoctoral Fellowship (23PRECHF1025586), and the NIDCR Interdisciplinary Training in Systems and Developmental Biology and Birth Defects (T32HD075735). Lastly, the differentiation schematic shown here was created using BioRender.com.

## Author Contributions

Conceptualization: C.Z.L., B.D.G.; Methodology: C.Z.L., B.D.G.; Validation: C.Z.L., B.D.G.; Formal Analysis: C.Z.L., B.D.G.; Investigation: C.Z.L., A.P., B.J.; Resources: B.D.G., A.J.S.; Data Curation: C.Z.L., B.J.; Writing—Original Draft: C.Z.L., B.D.G.; Visualization: C.Z.L.; Supervision: B.D.G., A.J.S.; Project Administration: B.D.G., A.J.S.; Funding Acquisition: C.Z.L., B.D.G.

## Resource Availability

### Lead Contact

Additional information or requests for reagents should be directed to the lead contact, Dr. Bruce Gelb.

### Materials availability

This study did not generate new unique reagents.

### Data and Code Availability

Data generated from the bulk RNA-sequencing were uploaded to GEO and can be accessed with accession number GSE226476.

## STAR METHODS

### EXPERIMENTAL MODEL AND SUBJECT DETAILS

#### Culture and maintenance of human iPSCs

Two wild-type human iPSC lines were utilized: WTC11 (XY, mono-allelic ACTN2-mEGFP, obtained from Coriell Institute #AICS-0075-085) and MSN02-4 (XX, generated from skin biopsy of a healthy 36-year-old at Icahn School of Medicine at Mount Sinai).^52, 53^ The iPSC lines were maintained in 6-well plates coated with a thin layer of Matrigel (6.6% v/v, Corning), diluted in DMEM (Gibco) and supplemented with penicillin/streptomycin (5%, ThermoFisher). iPSCs were maintained and fed daily with iPSC culture medium consisting of mTesR Plus (Stem Cell Technologies) supplemented with penicillin/streptomycin (1%, ThermoFisher). iPSCs were passaged every 3 to 4 days by first dissociating to single cells with Accutase (Innovative Cell Technologies, Inc) and then replating at a 1:10 ratio with iPSC culture medium, supplemented with Thiazovivin (2 μM, Selleckchem) for the first 24 h. iPSC cultures were incubated at 37 °C in a normoxic environment (5% CO_2_).

### METHOD DETAILS

#### Directed monolayer differentiation of iPSCs into endocardial & endothelial cells

Prior to the start of differentiation, iPSCs were passaged onto Matrigel coated 24-well plates at a ratio of 1:12. Day 0 (start of differentiation) was begun when iPSCs had reached ∼80-90% confluency. Each well was washed with DMEM (Gibco), followed by a media change into phase 1 media consisting of RPMI 1640 (Gibco) supplemented with B27 minus insulin supplement (0.5x, Gibco), penicillin/streptomycin (1%, ThermoFisher), GlutaMAX (2 mM, Gibco), ascorbic acid (50 μg/mL, Sigma), apo-transferrin (150 μg/mL, R&D Systems), and monothioglycerol (50 μg/mL, Sigma). CHIR-99021 (8 μM, Tocris) was added into phase 1 media for Day 0 of differentiation. For endocardial differentiation, media was changed after 48 h to phase 1 media supplemented with BMP10 (10 ng/mL, R&D Systems) and bFGF (50 ng/mL, R&D Systems). On Day 6, media was changed into phase 2 media consisting of RPMI 1640 (Gibco) supplemented with B27 supplement (0.5x, Gibco), penicillin/streptomycin (1%, ThermoFisher), Glutamax (2 mM, Gibco), ascorbic acid (50 μg/mL, Sigma), apo-transferrin (150 μg/mL, R&D Systems), and monothioglycerol (50 μg/mL, Sigma). Phase 2 media was further supplemented with BMP10 (10 ng/mL, R&D Systems) and bFGF (50 ng/mL, R&D Systems) for media changes between Days 6-14.

For endothelial differentiation, media change was performed after 48 h of CHIR-99021 (8 μM, Tocris) to phase 1 media supplemented with VEGF (100 ng/mL, R&D Systems) and bFGF (50 ng/ml, R&D Systems). On Day 6, media was changed to phase 2 media supplemented with VEGF (100 ng/mL, R&D Systems) and bFGF (50 ng/ml, R&D Systems) and maintained until Day 14. For both endocardial and endothelial conditions, media change was performed every other day and incubated at 37 °C in a hypoxic incubator (5% CO_2_, 5% O_2_, 90% N_2_).

#### Cell sorting and flow cytometry

For mesoderm analysis, cells undergoing either endocardial or endothelial differentiation were dissociated on Day 4 with Accutase (Innovative Cell Technologies, Inc) to a single-cell suspension. Cells were then resuspended in FACS staining buffer consisting of PBS^-/-^ (Gibco) supplemented with FBS (10%, Gibco), DNase (10 μg/mL, Millipore), and Thiazovivin (2 μM, Selleckchem) and labeled with anti-CD56 APC (1:100, BioLegend) and anti-CD13 PE (1:100, BioLegend) on ice for 1 h. Following washes with PBS^-/-^ (Gibco), cells were resuspended in FACS running buffer consisting of PBS^-/-^ (Gibco) supplemented with FBS (1%, Gibco), penicillin/streptomycin (2%, ThermoFisher), DNase (10 μg/mL, Millipore), thiazovivin (2 μM, Selleckchem), and DAPI (0.1 μg/mL, Sigma).

For endocardial and endothelial analysis, cells were dissociated with collagenase type II (0.6 mg/ml, Worthington) in HBSS (Corning) supplemented with HEPES (5 mM, Sigma) and incubated at 37 °C for 1 h. Dissociated cells were collected on ice with wash media consisting of DMEM (Gibco), BSA (0.05%, Gibco), DNAse (10 μg/mL, Millipore) and thiazovivin (2 μM, Selleckchem) and passed through a 40-μm cell strainer (Fisher Scientific). Enrichment of CD31^+^ endocardial cells or CD34^+^ endothelial cells to a purity greater than 95% was then performed by magnetic activated cell sorting (MACS, Miltenyi). Cells from the endocardial differentiation were incubated with human anti-CD31 microbeads (Miltenyi #130-091-935) according to manufacturer instructions in EasySep Buffer (Stem Cell Technologies) for 15 min on ice, followed by purification on an LS column. Similarly, cells from the endothelial differentiation were incubated with human anti-CD34 microbeads (Miltenyi #130-046-702) and purified on an LS column. Both endocardial and endothelial populations were stained with anti-CD31 APC (1:100, Invitrogen) and anti-CD34 PE-Cy7 (1:100, BioLegend) and analyzed via flow cytometry as described above.

Following EndMT, cells were dissociated with collagenase type II (0.6 mg/ml, Worthington) and collected as described above. VIC and VEC populations were separated with human CD31 MACS kit (Miltenyi #130-091-935) and purified on an LS column. Negative selection was performed to isolate CD31^-^ VICs, which are not magnetically bound to the LS column. Conversely, the cells bound to the LS column were then eluted to isolate CD31^+^ VECs. These populations were stained with anti-CD31 APC (1:100, Invitrogen) and anti-PDGFR*β* BV786 (1:100, BD Biosciences) and analyzed via flow cytometry as described above. All samples were analyzed on either the BD FACSCelesta, BD LSRFortessa, or BD LSR II flow cytometers.

#### EndMT induction of endocardial cells to generate VICs and VECs

On Day 14 of endocardial differentiation, endocardial cells were dissociated with collagenase type II (0.6 mg/ml, Worthington) for 1 h and enriched for a CD31^+^ endocardial population with CD31 MACS, as described above. Following purification, cell counts were determined with an automated cell counter (TC20, BioRad) and replated 3×10^5^ cells/well on fibronectin-coated 24-well plates in the EndMT condition consisting of phase 2 media supplemented with BMP2 (100 ng/ml, R&D Systems), TGF*β*2 (0.3 ng/mL, R&D Systems), and bFGF (10 ng/mL, R&D Systems). Thiazovivin (2 μM, Selleckchem) was also included for the first 16 h following plating. Fibronectin coating was performed with human fibronectin plasma (1:100, Sigma) diluted in PBS^-/-^ (Gibco) and incubated for at least 1 h at 37 °C. Plated cells were maintained in the EndMT condition for 6-7 days with media changes every other day and incubated at 37 °C in a hypoxic incubator (5% CO_2_, 5% O_2_, 90% N_2_).

#### Immunohistochemistry

Day 14 endocardial and endothelial cells were dissociated with collagenase type II (0.6 mg/ml, Worthington) and sorted with CD31-MACS and CD34-MACS, respectively, as described above. Endocardial and endothelial cells were plated on 12-chamber microscope slides (Ibidi) coated with Matrigel (6.6% v/v, Corning). Endocardial cells (1×10^5^) were plated with phase 2 media supplemented with BMP10 (10 ng/mL, R&D Systems) and bFGF (50 ng/mL, R&D Systems). Endothelial cells (1×10^5^) were plated with phase 2 media supplemented with VEGF (100 ng/mL, R&D Systems) and bFGF (50 ng/ml, R&D Systems).

Following EndMT, VICs and VECs were dissociated with collagenase type II (0.6 mg/ml, Worthington) and sorted with CD31-MACS for positive and negative selection, as described above. Isolated VICs and VECs were plated on 12-chamber microscopy slides (Ibidi) coated with human fibronectin plasma (1:100, Sigma). VICs were plated in fibroblast growth media (PromoCell) supplemented with penicillin/streptomycin (1%, ThermoFisher) and FBS (10%, Gibco). VECs were plated in MV2 endothelial media (PromoCell) supplemented with penicillin/streptomycin (1%, ThermoFisher).

All cells were incubated at 37 °C overnight in a hypoxic incubator (5% CO_2_, 5% O_2_, 90% N_2_). Cells were fixed with 4% PFA in PBS (ThermoFisher) for 10 min at room temperature, followed by permeabilization with 0.1% Triton X-100 in PBS for 10 min. Cells were washed with PBS and blocked for 1 h at room temperature with PBS containing BSA (1%, Sigma), glycine (22.52 mg/mL, BioRad), Tween-20 (0.1%, Sigma), and sodium azide (0.02%, Sigma). Incubation with primary antibodies was performed overnight at 4 °C in PBS containing BSA (1%, Sigma), Tween-20 (0.1%, Sigma), and sodium azide (0.02%, Sigma). Stained cells were then washed 3x with PBS for 10 min each and stained with secondary antibodies for 1 h at room temperature, followed by another wash step. Cell nuclei were stained with DAPI (0.25 μg/mL, Sigma) in PBS for 10 min at room temperature. Coverslips were then mounted onto the samples with Vectashield Plus Antifade Mounting Medium (Vector Laboratories). The following primary antibodies were used: anti-NKX2.5 (1:100, Cell Signaling Technology), anti-CD31 (1:100, Invitrogen), anti-NFATc1 (1:50, Invitrogen), anti-CDH5 (1:100, Cell Signaling Technology), anti-VIMENTIN (1:100, Invitrogen), anti-PERIOSTIN (1:100, Santa Cruz Biotechnology), anti-SOX9 (1:100, Sigma), and anti-*α*SMA (1:100, Abcam). The following secondary antibodies were used for detection: goat anti-rabbit IgG Alexa Fluor 488 (1:400, Invitrogen), goat anti-mouse IgG Alexa Flour 488 (1:400, Invitrogen), goat anti-rabbit IgG Alexa Flour 568 (1:400, Invitrogen), and goat anti-mouse IgG Alexa Flour 568 (1:400, Invitrogen). Detailed antibody information is described in the Key Resources Table. Stained cells were imaged on a Zeiss LSM780 confocal microscope.

#### Quantitative reverse transcriptase PCR

Total RNA from iPSC-derived populations was isolated using the Qiagen RNeasy Micro Kit. Briefly, samples were lysed in Buffer RLT containing *β***-**mercaptoethanol (Sigma) according to manufacturer’s instructions. Samples were incubated with proteinase K (Qiagen) at 55 °C for 10 min prior to RNA isolation on the RNeasy column. On-column DNase treatment was performed according to manufacturer instructions and total RNA was eluted in RNase-free H_2_O. RNA quality was determined with the Agilent 2100 Bioanalyzer and the RNA 6000 Nano chip. RIN numbers were confirmed to be >8.0 for all samples. cDNA was generated using the SuperScript IV VILO Master Mix (Invitrogen). qRT-PCR was performed on a ViiA 7 Real-Time PCR System (Applied Biosystems) using the PowerTrack SYBR Green Master Mix (ThermoFisher). All experiments were run in triplicates using the *ΔΔ*Ct method relative to a human reference cDNA (Takara) and the housekeeping gene TBP. Primer sequences are listed in Table S1.

#### Bulk RNA sequencing analysis

Total RNA from sorted iPSC-derived populations was isolated as described above and confirmed to have a RIN > 8.0. Both library prep and sequencing were performed by Novogene as a single batch to minimize possible technical confounders. Library preparation was performed using polyA selection without strand-specificity and sequenced with an Illumina NovaSeq instrument using paired-end 150bp reads at a depth of ∼20 million reads per sample. Raw sequencing data in FASTQ format were mapped to the human reference genome (NCBI build38/UCSC hg38) with 2-pass mapping using STAR v2.7.10a.^54^ Duplicate reads were marked and removed with Picard v2.26.10 (http://broadinstitute.github.io/picard/). Gene counts were then obtained with HTSeq v2.0.2.^55^ Count matrices were then analyzed in R with the DESeq2 package.^56^ To reduce variability in our dataset stemming from sex differences between the WTC11 and MSN02-4 iPSC lines, genes on the X and Y chromosomes were filtered out prior to differential analysis. Differential analysis was performed with DESeq2 using the likelihood ratio test. Heatmaps were generated using the R package ‘pheatmap’ on count data normalized with regularized log transformation. Volcano plots were generated using the R package ‘Enhanced Volcano’ and performed on count data following adaptive shrinkage estimation with ‘ashr’.^57^ Gene ontology analysis was performed with Enrichr using either the GO Biological Processes 2021 or BioPlanet 2019 annotations.^58^

#### Quantification and statistical analysis

All data are represented as mean ± standard error of mean (SEM). Biological replicates of each differentiation experiment are represented by the sample size (n). Sample sizes were not predetermined and due to the nature of the experiments, samples were not randomized or blinded from investigators. Statistical significance was determined using 2-way ANOVA ± multiple comparison analysis with Sidak, Tukey, or Dunnet’s post hoc test in GraphPad Prism 9 software. Results were marked as significant at the following cutoffs p < 0.05 (*), p < 0.01 (**), p < 0.001 (***), p < 0.0001 (****). All statistical tests and parameters are reported in the respective figures and figure legends.

## Figure Legends

### Supplemental Figure Legends

**Figure S1:**
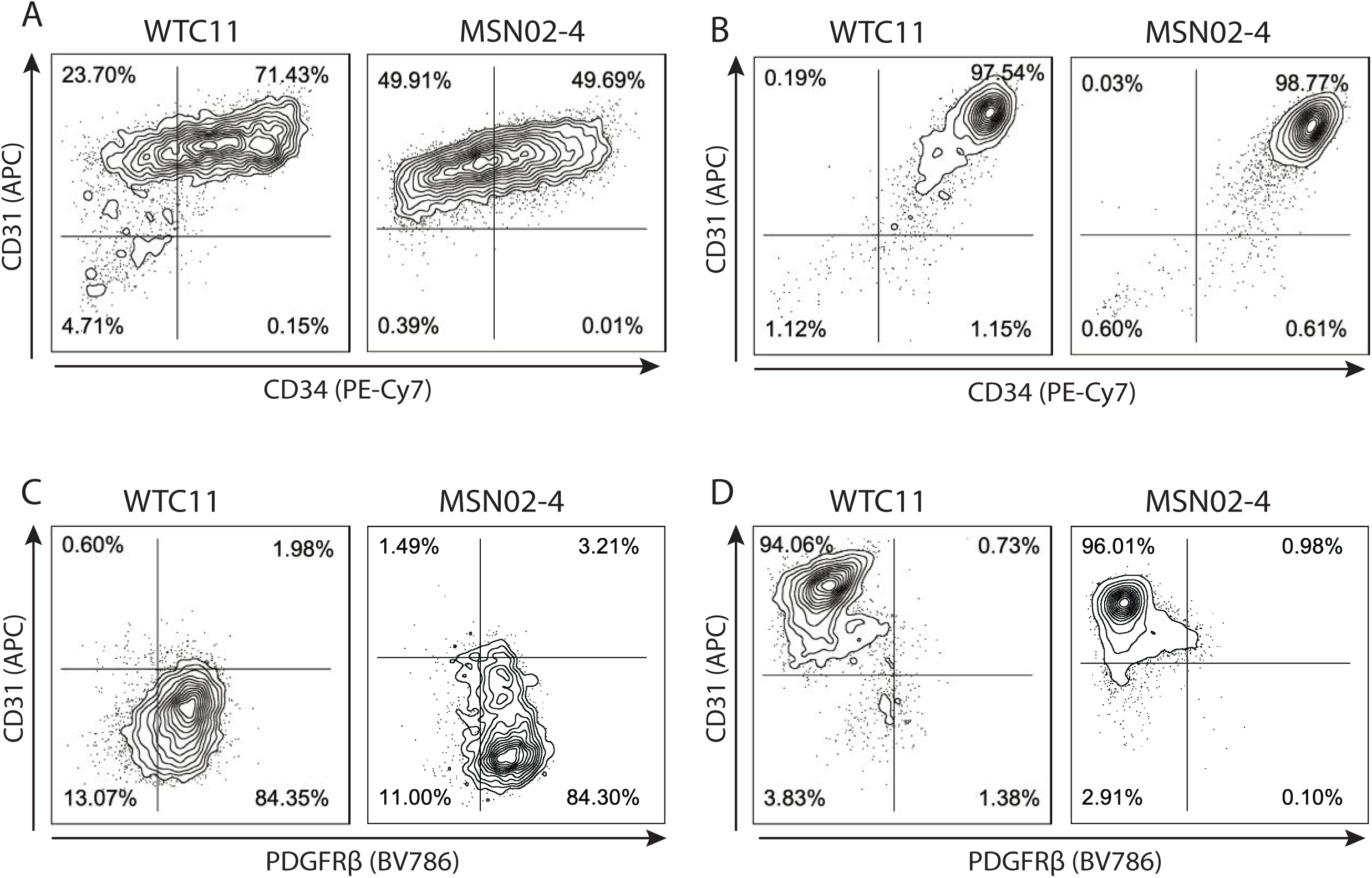
Validation of sort efficiencies following MACS. **(A)** Sort efficiency following CD31 MACS on Day 14 endocardial cells. **(B)** Sort efficiency following CD34 MACS on Day 14 endothelial cells. **(C)** Sort efficiency following negative selection with CD31 MACS for post-EndMT VICs. **(D)** Sort efficiency following positive selection with CD31 MACS for post-EndMT VECs.

**Figure S2:**
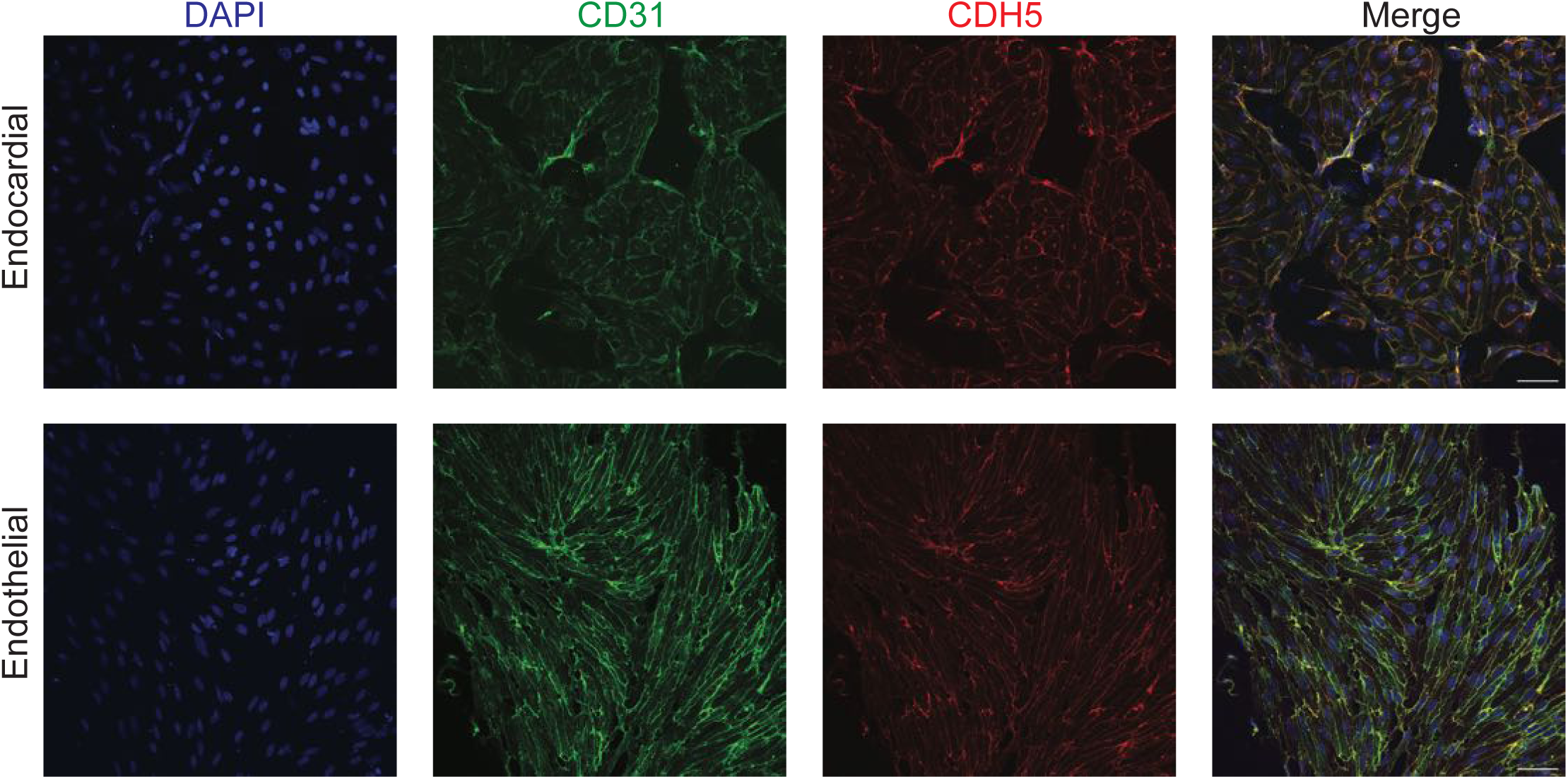
Confocal microscopy of CD31 and CDH5 of endocardial and endothelial cells. Endocardial and endothelial cells stained for CD31 (GFP) and CDH5 (RFP).

**Figure S3:**
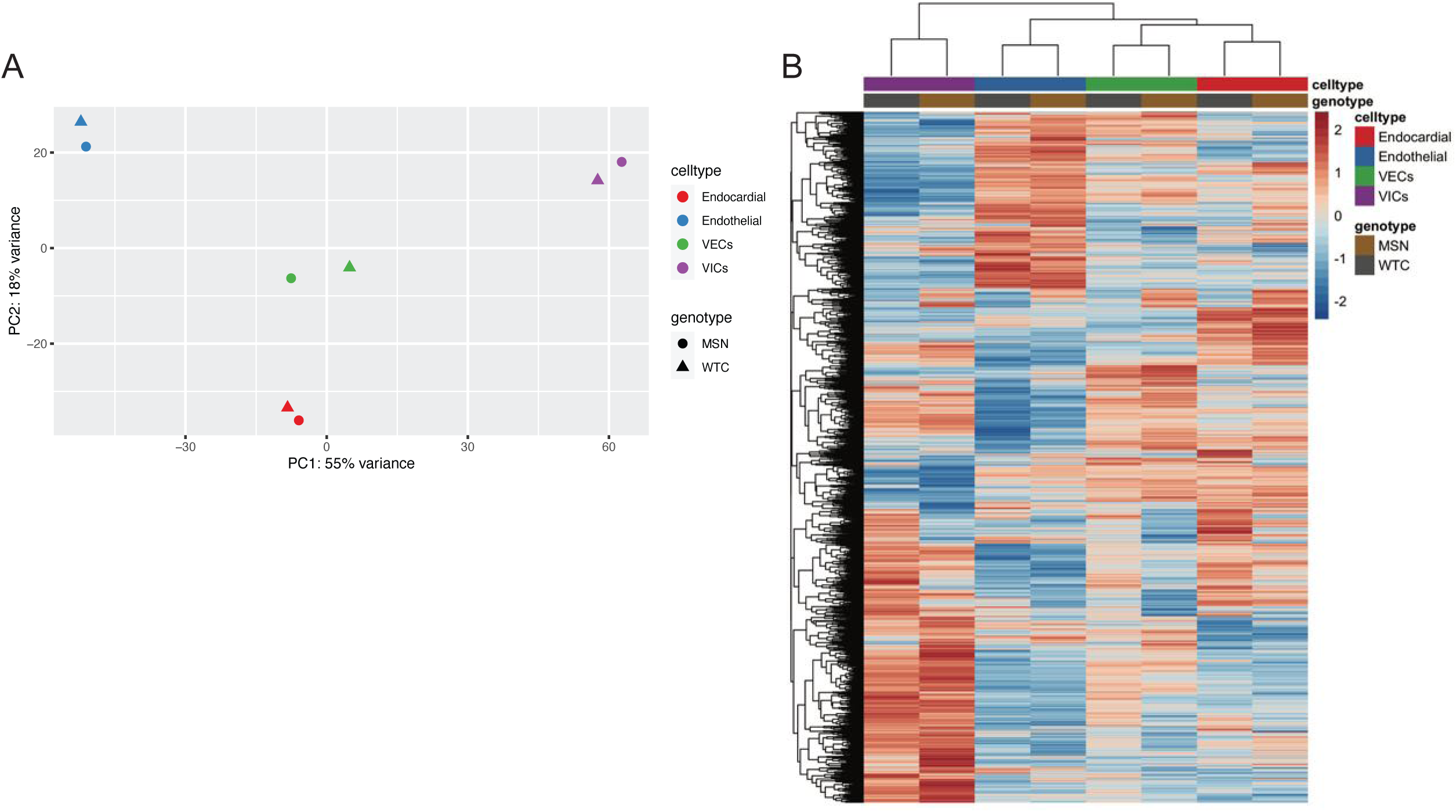
Bulk-RNAseq analysis on all cell populations. **(A)** PCA plot of each of the cell populations from both WTC11 and MSN02-4 iPSC lines. **(B)** Heatmap with unsupervised clustering on the top quartile of genes ranked by variance for all cell types.

**Figure S4:**
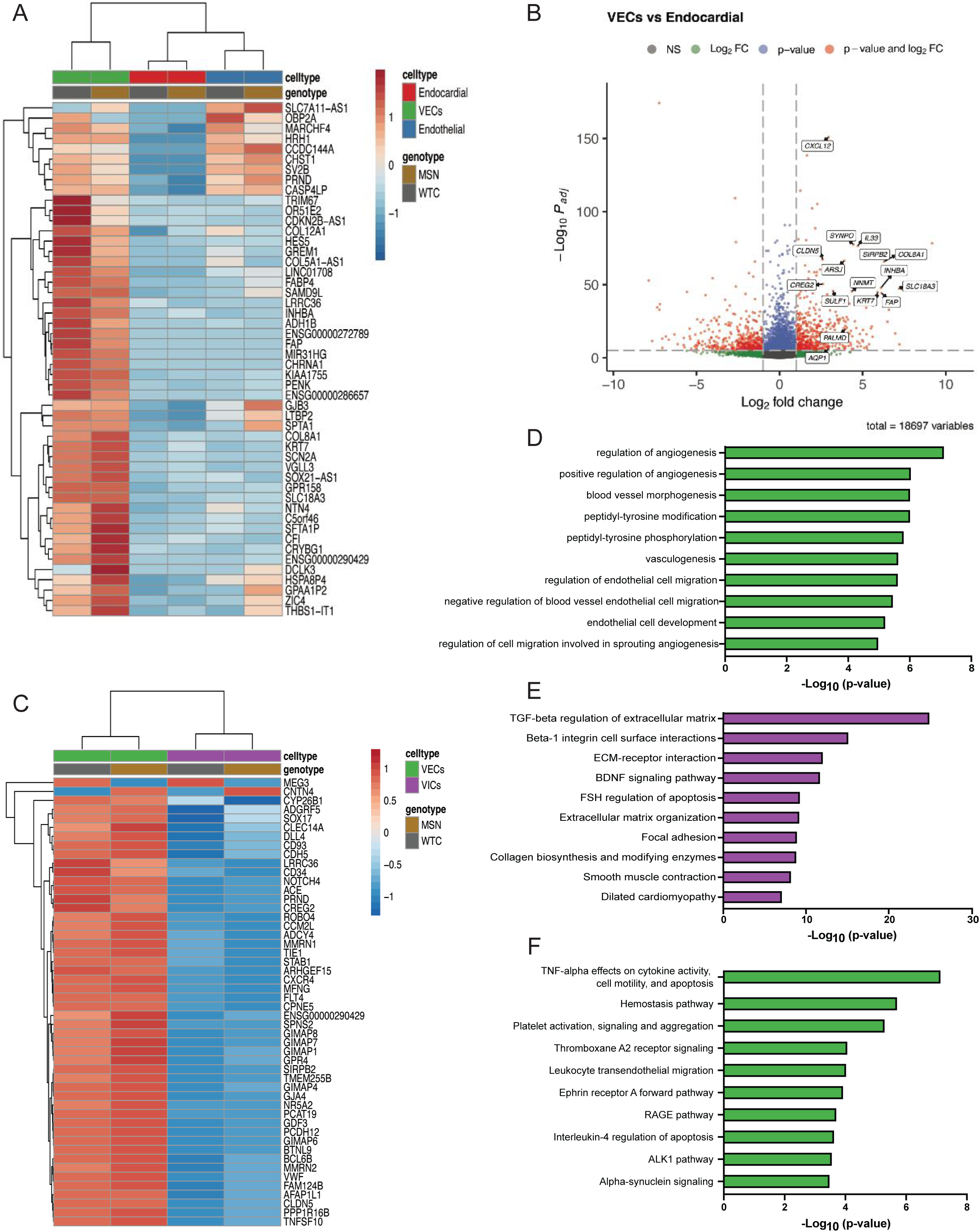
Bulk-RNAseq analysis of VECs. **(A)** Heatmap of the top 50 DEGs upregulated in VECs compared to endocardial and endothelial populations. **(B)** Volcano plot of DEGs between VEC and endocardial populations. **(C)** Heatmap of top 50 DEGs ranked by variance between VECs and VICs. **(D)** Top 10 biological process GO terms for upregulated DEGs in VECs relative to VICs. **(E)** Top 10 BioPlanet GO terms for DEGs upregulated in VICs relative to VECs. **(F)** Top 10 BioPlanet GO terms for DEGs upregulated in VECs relative to VICs.

**Table S1.**
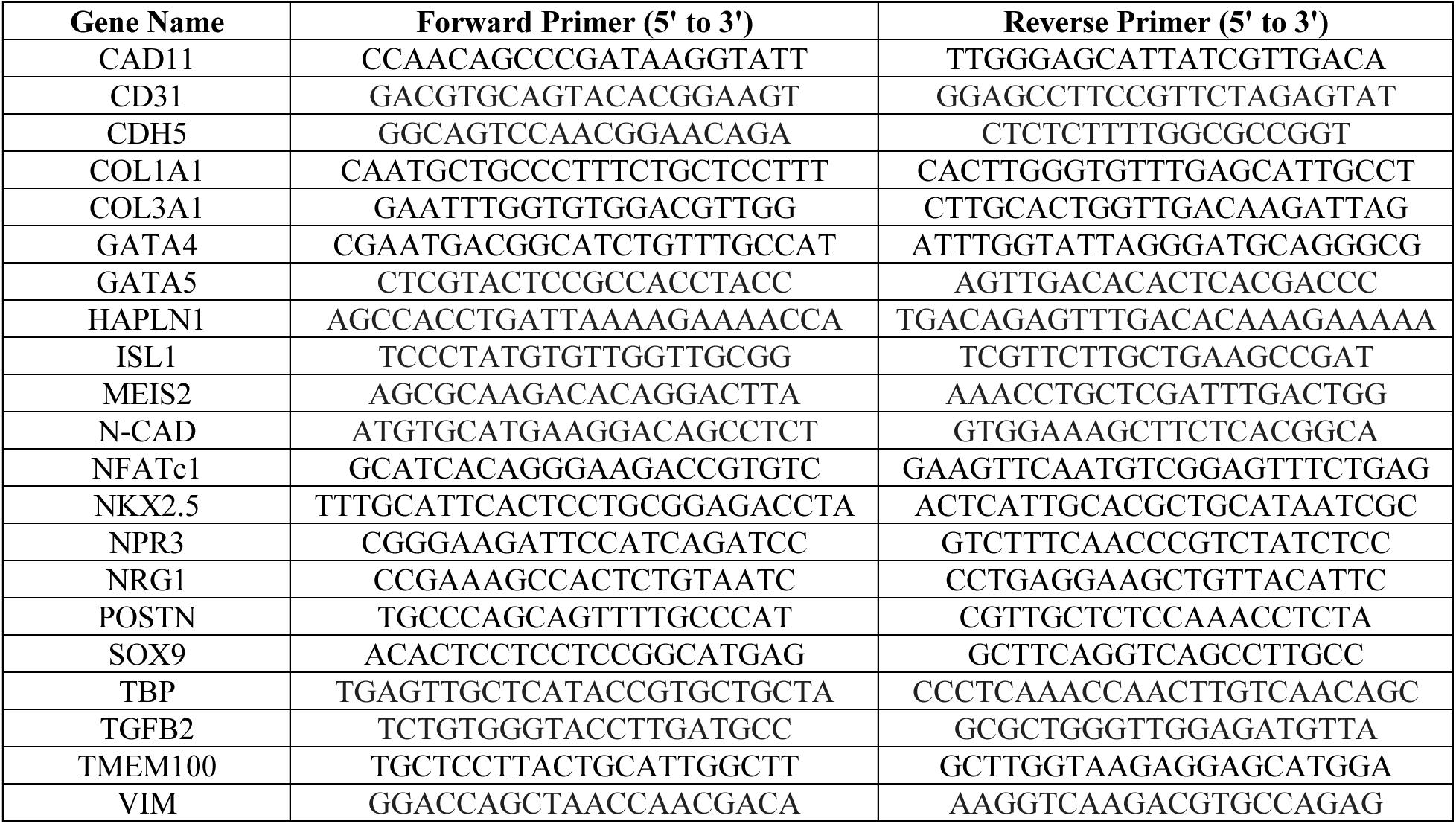
-List of Primers

Key resources table

Key resources table
| REAGENT or RESOURCE | SOURCE | IDENTIFIER |
| --- | --- | --- |
| **Antibodies** |  |  |
| Mouse monoclonal to CD31 (clone WM59), APC | ThermoFisher | Cat. #17-0319-42; RRID: AB_10852842 |
| Mouse monoclonal to CD31 (clone JC/70A) | ThermoFisher | Cat. #MA5-13188; RRID: AB_10982120 |
| Mouse monoclonal to NFATC1 (clone 7A6) | ThermoFisher | Cat. #MA3-024; RRID: AB_2236037 |
| Rabbit monoclonal to VIMENTIN (clone SP20) | ThermoFisher | Cat. #MA5-16409; RRID: AB_2537928 |
| Mouse monoclonal to CD34 (clone 561), PE-Cy7 | BioLegend | Cat. #343616; RRID: AB_2629726 |
| Mouse monoclonal to CD56 (clone 5.1H11), APC | BioLegend | Cat. #362504; RRID: AB_2563913 |
| Mouse monoclonal to CD13 (clone WM15), PE | BioLegend | Cat. #301704; RRID: AB_314180 |
| Rabbit monoclonal to NKX2.5 (clone E1Y8H) | Cell Signaling Technology | Cat. #8792; RRID: AB_2797667 |
| Rabbit polyclonal to VE-CADHERIN | Cell Signaling Technology | Cat. #2158; RRID: AB_2077970 |
| Mouse monoclonal to PERIOSTIN (clone F-10) | Santa Cruz Biotechnology | Cat. #sc-398631; RRID: AB_2934053 |
| Rabbit polyclonal to SOX9 | MilliporeSigma | Cat. #AB5535; RRID: AB_2239761 |
| Mouse monoclonal to αSMA (Clone 1A4) | Abcam | Cat. #ab7817; RRID: AB_262054 |
| Mouse monoclonal to PDGFRβ (clone 28D4), BV786 | BD Biosciences | Cat. #743038; RRID: AB_2741234 |
| Goat anti-Mouse IgG (H+L) Cross-Adsorbed Secondary Antibody, Alexa Fluor 488 | ThermoFisher | Cat. #A-1100; RRID: AB_2534069 |
| Goat anti-Rabbit IgG (H+L) Cross-Adsorbed Secondary Antibody, Alexa Fluor 488 | ThermoFisher | Cat. #A-11008; RRID: AB_143165 |
| Goat anti-Mouse IgG (H+L) Cross-Adsorbed Secondary Antibody, Alexa Fluor 568 | ThermoFisher | Cat. #A-11004; RRID: AB_2534072 |
| Goat anti-Rabbit IgG (H+L) Cross-Adsorbed Secondary Antibody, Alexa Fluor 568 | ThermoFisher | Cat. #A-11011; RRID: AB_143157 |
| **Chemicals, peptides, and recombinant proteins** |  |  |
| Recombinant Human BMP10 | R&D | Cat. #2926-BP |
| Recombinant Human bFGF | R&D | Cat. #233-FB |
| Recombinant Human VEGF-165 | R&D | Cat. #293-VE |
| Recombinant Human/Mouse/Rat BMP2 | R&D | Cat. #355-BEC |
| Recombinant Human TGF-beta 2 | R&D | Cat. #302-B2 |
| Human Apo-Transferrin | R&D | Cat. #3188-AT |
| Penicillin/streptomycin | ThermoFisher | Cat. #15140122 |
| GlutaMAX | ThermoFisher | Cat. #35050061 |
| Bovine Serum Albumin Fraction V (BSA) | ThermoFisher | Cat. #15260037 |
| Ascorbic Acid | MilliporeSigma | Cat. #A4544 |
| 2-Mercaptoethanol | MilliporeSigma | Cat. #M3148 |
| DAPI | MilliporeSigma | Cat. #9564 |
| Fetal Bovine Serum (FBS) | MilliporeSigma | Cat. #F4135 |
| Fibronectin Human Plasma | MilliporeSigma | Cat. #F0895 |
| Triton X-100 | MilliporeSigma | Cat. #93418 |
| Sodium Azide | MilliporeSigma | Cat. #S2002 |
| Tween-20 | MilliporeSigma | Cat. #P1379 |
| DNase I | MilliporeSigma | Cat. # 260913 |
| DMSO | MilliporeSigma | Cat. #D2650 |
| 4% Paraformaldehyde | ThermoFisher | Cat. # J19943K2 |
| CHIR 99021 | Tocris | Cat. #4423 |
| Collagenase Type II | Worthington | Cat. #LS004176 |
| Thiazovivin | Selleckchem | Cat. #S1459 |
| Matrigel, Growth Factor Reduced | Corning | Cat. #354230 |
| HEPES | Research Products International | Cat. #H75050 |
| Glycine | Bio-Rad | Cat. #1610718 |
| Proteinase K | Qiagen | Cat. #19131 |
| Accutase | Accutase | Cat. #AT104 |
| **Critical commercial assays** |  |  |
| Superscript IV VILO | ThermoFisher | Cat. #11756050 |
| PowerTrack SYBR Green | ThermoFisher | Cat. #A46012 |
| RNA 6000 Nano Kit | Agilent | Cat. # 5067-1511 |
| CD31 MicroBead Kit | Miltenyi Biotec | Cat. #130-091-935 |
| CD34 MicroBead Kit | Miltenyi Biotec | Cat. #130-046-702 |
| **Deposited data** |  |  |
| Raw & processed data from iPSC-derived populations | This study | GEO: GSE226476 |
| **Experimental models: Cell lines** |  |  |
| WTC11 (ACTN2-eGFP tagged, monoallelic) | Coriell Institute | Cat. #AICS-0075-085 |
| MSN02-4<sup>52,53</sup> | Gift from Dr. Dubois | N/A |
| **Oligonucleotides** |  |  |
| See Table S1 | This study | Table S1 |
| **Software and algorithms** |  |  |
| FCS Express 7 | De Novo Software | https://denovosoftware.com/ |
| GraphPad Prism | GraphPad | https://www.graphpad.com/ |
| R CRAN, Version 4.2.0 |  | https://cran.r-project.org/ |
| STAR v.2.7.10a<sup>54</sup> |  | https://github.com/alexdobin/STAR |
| Picard v2.26.10 |  | http://broadinstitute.github.io/picard/ |
| HTSeq v2.0.2<sup>55</sup> |  | https://htseq.readthedocs.io/en/master/ |
| DESeq2<sup>56</sup> |  | https://bioconductor.org/packages/release/bioc/html/DESeq2.html |
| Enrichr<sup>58</sup> |  | https://maayanlab.cloud/Enrichr/ |
| FIJI |  | https://imagej.net/software/fiji/ |
| **Other** |  |  |
| RPMI 1640 | ThermoFisher | Cat. #11875093 |
| B27 Supplement | ThermoFisher | Cat. #17504044 |
| B27 Supplement, Minus Insulin | ThermoFisher | Cat. #A1895601 |
| DMEM, high glucose | ThermoFisher | Cat. #11965092 |
| PBS, no calcium, no magnesium | ThermoFisher | Cat. #14190144 |
| 6-well clear flat bottom TC-treated culture plates | Fisher Scientific | Cat. #FB012927 |
| 24-well clear flat bottom TC-treated culture plates | Fisher Scientific | Cat. #FB012929 |
| 40um cell strainer | Fisher Scientific | Cat. #22363547 |
| Fibroblast Growth Medium, Kit | PromoCell | Cat. #C-23110 |
| Endothelial Cell Growth Medium MV2, Kit | PromoCell | Cat. #C-22121 |
| HBSS | Corning | Cat. #21022CV |
| EasySep Buffer | StemCell Technologies | Cat. #20144 |
| 12-well chamber slide | Ibidi | Cat. #81201 |
| Coverslips | Ibidi | Cat. #10811 |
| VECTASHIELD PLUS Antifade Mounting Medium | Vector Laboratories | Cat. #H-1900 |
| LS Column | Miltenyi Biotec | Cat. #130-042-401 |
| Cell Counting Slides | Bio-Rad | Cat. #1450015 |
| Trypan Blue | Bio-Rad | Cat. #1450021 |
| Human Reference cDNA | Takara | Cat. #636693 |

## References

1. Nkomo, V.T., Gardin, J.M., Skelton, T.N., Gottdiener, J.S., Scott, C.G., and Enriquez-Sarano, M. (2006). Burden of valvular heart diseases: a population-based study. Lancet 368, 1005–1011. 10.1016/s0140-6736(06)69208-8.

2. Combs, M.D., and Yutzey, K.E. (2009). Heart Valve Development: Regulatory Networks in Development and Disease. Circ Res 105, 408–421. 10.1161/circresaha.109.201566.

3. Saef, J.M., and Ghobrial, J. (2021). Valvular heart disease in congenital heart disease: a narrative review. Cardiovasc Diagnosis Ther. 10.21037/cdt-19-693-b.

4. Borer, J.S., and Sharma, A. (2015). Drug Therapy for Heart Valve Diseases. Circulation 132, 1038–1045. 10.1161/circulationaha.115.016006.

5. O’Donnell, A., and Yutzey, K.E. (2020). Mechanisms of heart valve development and disease. Development 147, dev183020. 10.1242/dev.183020.

6. Tao, G., Kotick, J.D., and Lincoln, J. (2012). Heart Valve Development, Maintenance, and Disease The Role of Endothelial Cells. Curr Top Dev Biol 100, 203–232. 10.1016/b978-0-12-387786-4.00006-3.

7. Ma, L., Lu, M.-F., Schwartz, R.J., and Martin, J.F. (2005). Bmp2 is essential for cardiac cushion epithelial-mesenchymal transition and myocardial patterning. Development 132, 5601– 5611. 10.1242/dev.02156.

8. Eisenberg, L.M., and Markwald, R.R. (1995). Molecular Regulation of Atrioventricular Valvuloseptal Morphogenesis. Circ Res 77, 1–6. 10.1161/01.res.77.1.1.

9. McCulley, D.J., Kang, J., Martin, J.F., and Black, B.L. (2008). BMP4 is required in the anterior heart field and its derivatives for endocardial cushion remodeling, outflow tract septation, and semilunar valve development. Dev Dynam 237, 3200–3209. 10.1002/dvdy.21743.

10. Romano, L.A., and Runyan, R.B. (2000). Slug is an Essential Target of TGFβ2 Signaling in the Developing Chicken Heart. Dev Biol 223, 91–102. 10.1006/dbio.2000.9750.

11. Timmerman, L.A., Grego-Bessa, J., Raya, A., Bertrán, E., Pérez-Pomares, J.M., Díez, J., Aranda, S., Palomo, S., McCormick, F., Izpisúa-Belmonte, J.C., et al. (2004). Notch promotes epithelial-mesenchymal transition during cardiac development and oncogenic transformation. Gene Dev 18, 99–115. 10.1101/gad.276304.

12. MacGrogan, D., D’Amato, G., Travisano, S., Martinez-Poveda, B., Luxán, G., Monte-Nieto, G. del, Papoutsi, T., Sbroggio, M., Bou, V., Arco, P.G., et al. (2016). Sequential Ligand-Dependent Notch Signaling Activation Regulates Valve Primordium Formation and Morphogenesis. Circ Res 118, 1480–1497. 10.1161/circresaha.115.308077.

13. Liebner, S., Cattelino, A., Gallini, R., Rudini, N., Iurlaro, M., Piccolo, S., and Dejana, E. (2004). Beta-catenin is required for endothelial-mesenchymal transformation during heart cushion development in the mouse. J Cell Biology 166, 359–367. 10.1083/jcb.200403050.

14. Zhang, H., Gise, A. von, Liu, Q., Hu, T., Tian, X., He, L., Pu, W., Huang, X., He, L., Cai, C.-L., et al. (2014). Yap1 Is Required for Endothelial to Mesenchymal Transition of the Atrioventricular Cushion. J Biol Chem 289, 18681–18692. 10.1074/jbc.m114.554584.

15. Hinton, R.B., and Yutzey, K.E. (2011). Heart Valve Structure and Function in Development and Disease. Physiology 73, 29–46. 10.1146/annurev-physiol-012110-142145.

16. Kodigepalli, K.M., Thatcher, K., West, T., Howsmon, D.P., Schoen, F.J., Sacks, M.S., Breuer, C.K., and Lincoln, J. (2020). Biology and Biomechanics of the Heart Valve Extracellular Matrix. J Cardiovasc Dev Dis 7, 57. 10.3390/jcdd7040057.

17. Liu, A.C., Joag, V.R., and Gotlieb, A.I. (2007). The Emerging Role of Valve Interstitial Cell Phenotypes in Regulating Heart Valve Pathobiology. Am J Pathology 171, 1407–1418. 10.2353/ajpath.2007.070251.

18. Nakamura, T., Colbert, M.C., and Robbins, J. (2006). Neural Crest Cells Retain Multipotential Characteristics in the Developing Valves and Label the Cardiac Conduction System. Circ Res 98, 1547–1554. 10.1161/01.res.0000227505.19472.69.

19. Jain, R., Engleka, K.A., Rentschler, S.L., Manderfield, L.J., Li, L., Yuan, L., and Epstein, J.A. (2011). Cardiac neural crest orchestrates remodeling and functional maturation of mouse semilunar valves. J Clin Invest 121, 422–430. 10.1172/jci44244.

20. Wessels, A., Hoff, M.J.B. van den, Adamo, R.F., Phelps, A.L., Lockhart, M.M., Sauls, K., Briggs, L.E., Norris, R.A., Wijk, B. van, Perez-Pomares, J.M., et al. (2012). Epicardially derived fibroblasts preferentially contribute to the parietal leaflets of the atrioventricular valves in the murine heart. Dev Biol 366, 111–124. 10.1016/j.ydbio.2012.04.020.

21. Lockhart, M.M., Boukens, B.J.D., Phelps, A.L., Brown, C.-L.M., Toomer, K.A., Burns, T.A., Mukherjee, R.D., Norris, R.A., Trusk, T.C., Hoff, M.J.B. van den, et al. (2014). Alk3 mediated Bmp signaling controls the contribution of epicardially derived cells to the tissues of the atrioventricular junction. Dev Biol 396, 8–18. 10.1016/j.ydbio.2014.09.031.

22. Liu, K., Yu, W., Tang, M., Tang, J., Liu, X., Liu, Q., Li, Y., He, L., Zhang, L., Evans, S.M., et al. (2018). A dual genetic tracing system identifies diverse and dynamic origins of cardiac valve mesenchyme. Development 145, dev167775. 10.1242/dev.167775.

23. Kim, A.J., Xu, N., and Yutzey, K.E. (2020). Macrophage lineages in heart valve development and disease. Cardiovasc Res 117, 663–673. 10.1093/cvr/cvaa062.

24. Shigeta, A., Huang, V., Zuo, J., Besada, R., Nakashima, Y., Lu, Y., Ding, Y., Pellegrini, M., Kulkarni, R.P., Hsiai, T., et al. (2019). Endocardially Derived Macrophages Are Essential for Valvular Remodeling. Dev Cell 48, 617–630.e3. 10.1016/j.devcel.2019.01.021.

25. Neri, T., Hiriart, E., Vliet, P.P. van, Faure, E., Norris, R.A., Farhat, B., Jagla, B., Lefrancois, J., Sugi, Y., Moore-Morris, T., et al. (2019). Human pre-valvular endocardial cells derived from pluripotent stem cells recapitulate cardiac pathophysiological valvulogenesis. Nature Communications 10, 1929. 10.1038/s41467-019-09459-5.

26. Palpant, N.J., Pabon, L., Friedman, C.E., Roberts, M., Hadland, B., Zaunbrecher, R.J., Bernstein, I., Zheng, Y., and Murry, C.E. (2016). Generating high-purity cardiac and endothelial derivatives from patterned mesoderm using human pluripotent stem cells. Nat Protoc 12, 15–31. 10.1038/nprot.2016.153.

27. Palpant, N.J., Pabon, L., Roberts, M., Hadland, B., Jones, D., Jones, C., Moon, R.T., Ruzzo, W.L., Bernstein, I., Zheng, Y., et al. (2015). Inhibition of β-catenin signaling respecifies anterior-like endothelium into beating human cardiomyocytes. Development 142, 3198–3209. 10.1242/dev.117010.

28. Cheng, L., Xie, M., Qiao, W., Song, Y., Zhang, Y., Geng, Y., Xu, W., Wang, L., Wang, Z., Huang, K., et al. (2021). Generation and characterization of cardiac valve endothelial-like cells from human pluripotent stem cells. Commun Biology 4, 1039. 10.1038/s42003-021-02571-7.

29. Bao, X., Bhute, V.J., Han, T., Qian, T., Lian, X., and Palecek, S.P. (2017). Human pluripotent stem cell-derived epicardial progenitors can differentiate to endocardial-like endothelial cells. Bioeng Amp Transl Medicine 2, 191–201. 10.1002/btm2.10062.

30. Mikryukov, A.A., Mazine, A., Wei, B., Yang, D., Miao, Y., Gu, M., and Keller, G.M. (2021). BMP10 Signaling Promotes the Development of Endocardial Cells from Human Pluripotent Stem Cell-Derived Cardiovascular Progenitors. Cell Stem Cell 28, 1–16. 10.1016/j.stem.2020.10.003.

31. Theunissen, T.W., Powell, B.E., Wang, H., Mitalipova, M., Faddah, D.A., Reddy, J., Fan, Z.P., Maetzel, D., Ganz, K., Shi, L., et al. (2014). Systematic Identification of Culture Conditions for Induction and Maintenance of Naive Human Pluripotency. Cell Stem Cell 15, 471–487. 10.1016/j.stem.2014.07.002.

32. Yue, X.-S., Fujishiro, M., Nishioka, C., Arai, T., Takahashi, E., Gong, J.-S., Akaike, T., and Ito, Y. (2012). Feeder Cells Support the Culture of Induced Pluripotent Stem Cells Even after Chemical Fixation. Plos One 7, e32707. 10.1371/journal.pone.0032707.

33. Wan, H., Fu, R., Tong, M., Wang, Y., Wang, L., Wang, S., Zhang, Y., Li, W., Wang, X., and Feng, G. (2022). Influence of feeder cells on transcriptomic analysis of pluripotent stem cells. Cell Proliferat 55, e13189. 10.1111/cpr.13189.

34. Lian, X., Hsiao, C., Wilson, G., Zhu, K., Hazeltine, L.B., Azarin, S.M., Raval, K.K., Zhang, J., Kamp, T.J., and Palecek, S.P. (2012). Robust cardiomyocyte differentiation from human pluripotent stem cells via temporal modulation of canonical Wnt signaling. Proc National Acad Sci 109, E1848–E1857. 10.1073/pnas.1200250109.

35. Lian, X., Zhang, J., Azarin, S.M., Zhu, K., Hazeltine, L.B., Bao, X., Hsiao, C., Kamp, T.J., and Palecek, S.P. (2013). Directed cardiomyocyte differentiation from human pluripotent stem cells by modulating Wnt/β-catenin signaling under fully defined conditions. Nat Protoc 8, 162– 175. 10.1038/nprot.2012.150.

36. Sriram, G., Tan, J.Y., Islam, I., Rufaihah, A.J., and Cao, T. (2015). Efficient differentiation of human embryonic stem cells to arterial and venous endothelial cells under feeder- and serum-free conditions. Stem Cell Res Ther 6, 261. 10.1186/s13287-015-0260-5.

37. Tan, J.Y., Sriram, G., Rufaihah, A.J., Neoh, K.G., and Cao, T. (2013). Efficient Derivation of Lateral Plate and Paraxial Mesoderm Subtypes from Human Embryonic Stem Cells Through GSKi-Mediated Differentiation. Stem Cells Dev 22, 1893–1906. 10.1089/scd.2012.0590.

38. Batalov, I., and Feinberg, A.W. (2015). Differentiation of Cardiomyocytes from Human Pluripotent Stem Cells Using Monolayer Culture. Biomark Insights 10s1, BMI.S20050. 10.4137/bmi.s20050.

39. Evseenko, D., Zhu, Y., Schenke-Layland, K., Kuo, J., Latour, B., Ge, S., Scholes, J., Dravid, G., Li, X., MacLellan, W.R., et al. (2010). Mapping the first stages of mesoderm commitment during differentiation of human embryonic stem cells. Proc National Acad Sci 107, 13742– 13747. 10.1073/pnas.1002077107.

40. Skelton, R.J.P., Brady, B., Khoja, S., Sahoo, D., Engel, J., Arasaratnam, D., Saleh, K.K., Abilez, O.J., Zhao, P., Stanley, E.G., et al. (2016). CD13 and ROR2 Permit Isolation of Highly Enriched Cardiac Mesoderm from Differentiating Human Embryonic Stem Cells. Stem Cell Rep 6, 95–108. 10.1016/j.stemcr.2015.11.006.

41. Pompa, J.L. de la, Timmerman, L.A., Takimoto, H., Yoshida, H., Elia, A.J., Samper, E., Potter, J., Wakeham, A., Marengere, L., Langille, B.L., et al. (1998). Role of the NF-ATc transcription factor in morphogenesis of cardiac valves and septum. Nature 392, 182–186. 10.1038/32419.

42. Ranger, A.M., Grusby, M.J., Hodge, M.R., Gravallese, E.M., Brousse, F.C. de la, Hoey, T., Mickanin, C., Baldwin, H.S., and Glimcher, L.H. (1998). The transcription factor NF-ATc is essential for cardiac valve formation. Nature 392, 186–190. 10.1038/32426.

43. Monaghan, M.G., Linneweh, M., Liebscher, S., Handel, B.V., Layland, S.L., and Schenke-Layland, K. (2016). Endocardial-to-mesenchymal transformation and mesenchymal cell colonization at the onset of human cardiac valve development. Dev Camb Engl 143, 473–482. 10.1242/dev.133843.

44. Suehiro, J., Kanki, Y., Makihara, C., Schadler, K., Miura, M., Manabe, Y., Aburatani, H., Kodama, T., and Minami, T. (2014). Genome-wide Approaches Reveal Functional Vascular Endothelial Growth Factor (VEGF)-inducible Nuclear Factor of Activated T Cells (NFAT) c1 Binding to Angiogenesis-related Genes in the Endothelium. J Biol Chem 289, 29044–29059. 10.1074/jbc.m114.555235.

45. Kulkarni, R.M., Greenberg, J.M., and Akeson, A.L. (2009). NFATc1 regulates lymphatic endothelial development. Mech Develop 126, 350–365. 10.1016/j.mod.2009.02.003.

46. Ridge, L.A., Kewbank, D., Schütz, D., Stumm, R., Scambler, P.J., and Ivins, S. (2021). Dual role for CXCL12 signaling in semilunar valve development. Cell Reports 36, 109610. 10.1016/j.celrep.2021.109610.

47. Wu, S., Kumar, V., Xiao, P., Kuss, M., Lim, J.Y., Guda, C., Butcher, J., and Duan, B. (2020). Age related extracellular matrix and interstitial cell phenotype in pulmonary valves. Sci Rep 10, 21338. 10.1038/s41598-020-78507-8.

48. Lotto, J., Cullum, R., Drissler, S., Arostegui, M., Garside, V., Fuglerud, B., Thakur, A., Underhill, T., and Hoodless, P. (2022). Single-cell transcriptomics reveals diversity during heart valve epithelial-to-mesenchymal transitions. Research Square. 10.21203/rs.3.rs-1647065/v1.

49. Hori, N., Okada, K., Takakura, Y., Takano, H., Yamaguchi, N., and Yamaguchi, N. (2020). Vestigial-like family member 3 (VGLL3), a cofactor for TEAD transcription factors, promotes cancer cell proliferation by activating the Hippo pathway. J Biol Chem 295, 8798–8807. 10.1074/jbc.ra120.012781.

50. Yu, W., Ma, X., Xu, J., Heumüller, A.W., Fei, Z., Feng, X., Wang, X., Liu, K., Li, J., Cui, G., et al. (2019). VGLL4 plays a critical role in heart valve development and homeostasis. Plos Genet 15, e1007977. 10.1371/journal.pgen.1007977.

51. Wang, M., Lin, B.Y., Sun, S., Dai, C., Long, F., and Butcher, J.T. (2022). Shear and hydrostatic stress regulate fetal heart valve remodeling through YAP-mediated mechanotransduction. bioRxiv. 10.1101/2022.11.24.517814.

52. Schaniel, C., Dhanan, P., Hu, B., Xiong, Y., Raghunandan, T., Gonzalez, D.M., Dariolli, R., D’Souza, S.L., Yadaw, A.S., Hansen, J., et al. (2021). A library of induced pluripotent stem cells from clinically well-characterized, diverse healthy human individuals. Stem Cell Rep 16, 3036– 3049. 10.1016/j.stemcr.2021.10.005.

53. Wickramasinghe, N.M., Sachs, D., Shewale, B., Gonzalez, D.M., Dhanan-Krishnan, P., Torre, D., LaMarca, E., Raimo, S., Dariolli, R., Serasinghe, M.N., et al. (2022). PPARdelta activation induces metabolic and contractile maturation of human pluripotent stem cell-derived cardiomyocytes. Cell Stem Cell 29, 559–576.e7. 10.1016/j.stem.2022.02.011.

54. Dobin, A., Davis, C.A., Schlesinger, F., Drenkow, J., Zaleski, C., Jha, S., Batut, P., Chaisson, M., and Gingeras, T.R. (2013). STAR: ultrafast universal RNA-seq aligner. Bioinformatics 29, 15–21. 10.1093/bioinformatics/bts635.

55. Putri, G.H., Anders, S., Pyl, P.T., Pimanda, J.E., and Zanini, F. (2022). Analysing high-throughput sequencing data in Python with HTSeq 2.0. Bioinformatics 38, 2943–2945. 10.1093/bioinformatics/btac166.

56. Love, M.I., Huber, W., and Anders, S. (2014). Moderated estimation of fold change and dispersion for RNA-seq data with DESeq2. Genome Biol 15, 550. 10.1186/s13059-014-0550-8.

57. Stephens, M. (2016). False discovery rates: a new deal. Biostatistics 18, 275–294. 10.1093/biostatistics/kxw041.

58. Chen, E.Y., Tan, C.M., Kou, Y., Duan, Q., Wang, Z., Meirelles, G.V., Clark, N.R., and Ma’ayan, A. (2013). Enrichr: interactive and collaborative HTML5 gene list enrichment analysis tool. Bmc Bioinformatics 14, 128. 10.1186/1471-2105-14-128.

